# CA3 hippocampal synaptic plasticity supports ripple physiology during memory consolidation

**DOI:** 10.1101/2023.03.28.534509

**Authors:** Hajer El Oussini, Chun-Lei Zhang, Urielle François, Cecilia Castelli, Aurélie Lampin-Saint-Amaux, Marilyn Lepleux, Pablo Molle, Legeolas Velez, Cyril Dejean, Frédéric Lanore, Cyril Herry, Daniel Choquet, Yann Humeau

## Abstract

Consolidation of recent memory depends on hippocampal activities during resting periods that immediately follows the memory encoding. There, Slow Save Sleep phases appear as privileged periods for memory consolidation as hosting the ripple activities, which are fast oscillations generated within the hippocampus whose inactivation leads to memory impairment. If a strong correlation exists between these replays of recent experience and the persistence of behavioural adaptations, the mobilisation, the localization and the importance of synaptic plasticity events in this process is largely unknown. To question this issue, we used cell-surface AMPAR immobilisation to block post-synaptic LTP within the hippocampal region at various steps of the memory process. 1- Our results show that hippocampal synaptic plasticity is engaged during the consolidation but is dispensable during the encoding or recall of a working memory based spatial memory task. 2- Blockade of plasticity during sleep leads to apparent forgetting of the encoded rule. 3- *In vivo* recordings of ripple activities during resting periods show a strong impact of AMPAR immobilization solely, prominent when a rule has been recently encoded. 4- *In situ* examination of the interplay between AMPAR mobility, hippocampal plasticity and spontaneous ripple activities pointed that post-synaptic plasticity at CA3-CA3 recurrent synapses support ripple generation. As crucial results were reproduced using another AMPARM blockade strategy, we propose that after rule encoding, post-synaptic AMPAR mobility at CA3 recurrent synapses support the generation of ripples necessary for rule consolidation.

## Introduction

The importance of activity-dependent synaptic plasticity (SP) in the memorization process is generally admitted. This statement is supported by multiple studies reporting the behavioural impact of pharmacological treatments targeting SP-related molecular mechanisms, SP-related genetic inactivation, or more recently molecular approaches specifically affecting long term potentiation^1–3^. The examination of the physiological consequences of these inactivation strategies in living animals generally point to alterations of behavioural performance at testing time ^2, 3^, an impact that however depends on the type of memory and the extent of molecular manipulations onto one or more implicated brain regions ^2^. Recent reports also challenged the link existing between neurobiological consequences of synaptic plasticity blockade and animal performances. For example, Kaganovsky and colleagues recently proposed that blocking synaptic plasticity in the CA1 region of the hippocampus did not impact animal performance in a variety of behavioural tests, even though some cellular and network proxies for learning were affected in the hippocampus ^2^. This may result from functional redundancy between brain regions to achieve cognitive robustness, but it also certainly complexifies the interpretation of the impact of these SP interference methods on memory.

One intriguing possibility would be that the process of memory that includes multiple steps – encoding, consolidation and retrieval – would correspond to sequential steps of synaptic plasticity formation and maintenance^4, 5^. There, Hebbian synaptic tagging would be the immediate response to coincident neuronal activities supporting rapid adaptation of animal behaviour in response to new situations. Then synaptic capture, necessary for the maintenance of the plasticity, would occur during quiescent - awake or sleeping - states, allowing consolidation of a memory that can be retrieved afterwards. Along this line, the importance of hippocampal ripples appears as central.

Ripples intrinsically contain both the prerequisite for synaptic capture, namely a capacity in replaying behaviourally relevant spatial sequences encoded during the awake state ^6^, but also to allow broadcasting of these information enabling expression of plasticity related molecules important for synaptic capture ^4^. Recent findings confirmed that ripple content depends on recently acquired memories^7^, reactivating neuronal ensembles in cortices, such as those implicated in the running of specific rules^8^.

Physiologically, hippocampal ripples are short network oscillations at 180-250 Hz corresponding to synchronized neuronal activation generating synaptic waves that can be evidenced if recording in the dendritic fields – *i.e. stratum radiatum* - of hippocampal CA3 and CA1 regions ^6^. Interestingly, ripples can reach cortical regions - through direct or indirect projections - where they synchronize with other sleep related oscillations, such as spindle and delta wav^6^e,sa process that is reinforced by newly encoded learning ^9^. In the hippocampus, ripples are generated in structures – such as in hippocampal CA3 region – rich in recurrent connectivity and depends both on excitatory and inhibitory local inputs that constitute the feedback loops necessary for fast oscillations buildi^1^n^0^g. As such, impairing ^11^ or prolongating ^7^ ripples can be achieved by manipulating specific interneuron populations .

Even if *in situ* preparations do allow spontaneous oscillations that share numerous characteristics with ripples recorded *in vivo*^10, 12, 13^ , the relation between synaptic plasticity and ripple physiology has not yet been explored specifically *in situ* by using methods avoiding confounding factors such as effects on basal glutamatergic transmission ^1^. We here used two alternative strategies to abolish synaptic plasticity depending on AMPAR mobility (AMPARM): the first, based on multivalent IGGs directed against the GluA2 subunit of AMPAR has been proved to be efficient in blocking LTP at Schaeffer collaterals to CA1 synapses ^14^. Similarly, blocking AMPARM using biotinylation of GluA2 subunit and presence of tetravalent neutravidin in the external medium did not affect basal transmission but led to a complete absence of LTP ^15^. We thus used these strategies in the dorsal hippocampus to explore the link between synaptic plasticity, ripple physiology and learning and memory processing.

## Results

### AMPAR immobilization in the dorsal hippocampus impairs memory consolidation

Based on our previous reports showing that AMPAR immobilization at neuronal surface was efficiently blocking post-synaptic expression of hippocampal long-term potentiation, we used intra- hippocampal infusions of AMPAR cross-linkers to test for the implication of synaptic plasticity in the memorization process of a spatial task. For this, mice were cannulated bilaterally above the dorsal hippocampus, and trained for working memory-based delayed spatial alternation task (DSA ^16^). In this task, food deprived mice are learned to find food rewards in a Y-maze according to a simple rule: reward location is alternating between right/left ending arms ( **Figure 1a**). A delay of 30 seconds between consecutive runs is imposed, forcing the animals to remember the previous location before engaging in the following run. In control conditions, a training day of 4 sessions - about 40 trials or reward positions - is sufficient for the animals to decrease its number of errors and reach its maximal performance which is maintained the following days ( **Figure 1d**). To mediate AMPAR immobilization in the dorsal hippocampus, we performed bilateral, intra-cerebral injections of AMPAR cross-linkers (anti-GluA2 IgGs) or their controls (anti GluA2 monovalent Fabs) at key timings of the learning process ( **Figure 1b-c**) : I m m e d i a t e l y b e f o r e t h e f i r s t l e a r n i n g s e s s i o n o f d a y immediately after the end of the first training day (Pre-rest), and immediately before the first session of day 2 (Pre-test). Our aim was to test the importance of hippocampal AMPARM-dependent plasticity in the encoding, the consolidation and the recall of DSA rule respectively. Collectively, results pointed to an impact of AMPAR cross-linking onto memory consolidation: Indeed, pre-learning injections of AMPAR cross-linkers did not impact animal performance on day 1 ( **Figure 1d and 1e _left_**), but rather on the following day, choices returning to random level ( **Figure 1d _left_ and 1e_right_**). A similar effect was observed when injections were performed immediately after session#4 (**Figure 1d_middle_, and 1f**), but not if done before the test performed in day2 (before session session#5, **Figure 1d _right_ and 1g**). This last experiment indicates that memory retrieval was not impacted by AMPAR cross-linking, but more importantly that crucial AMPARM-dependent process occurs during resting period that support memory consolidation.

**Figure 1:**
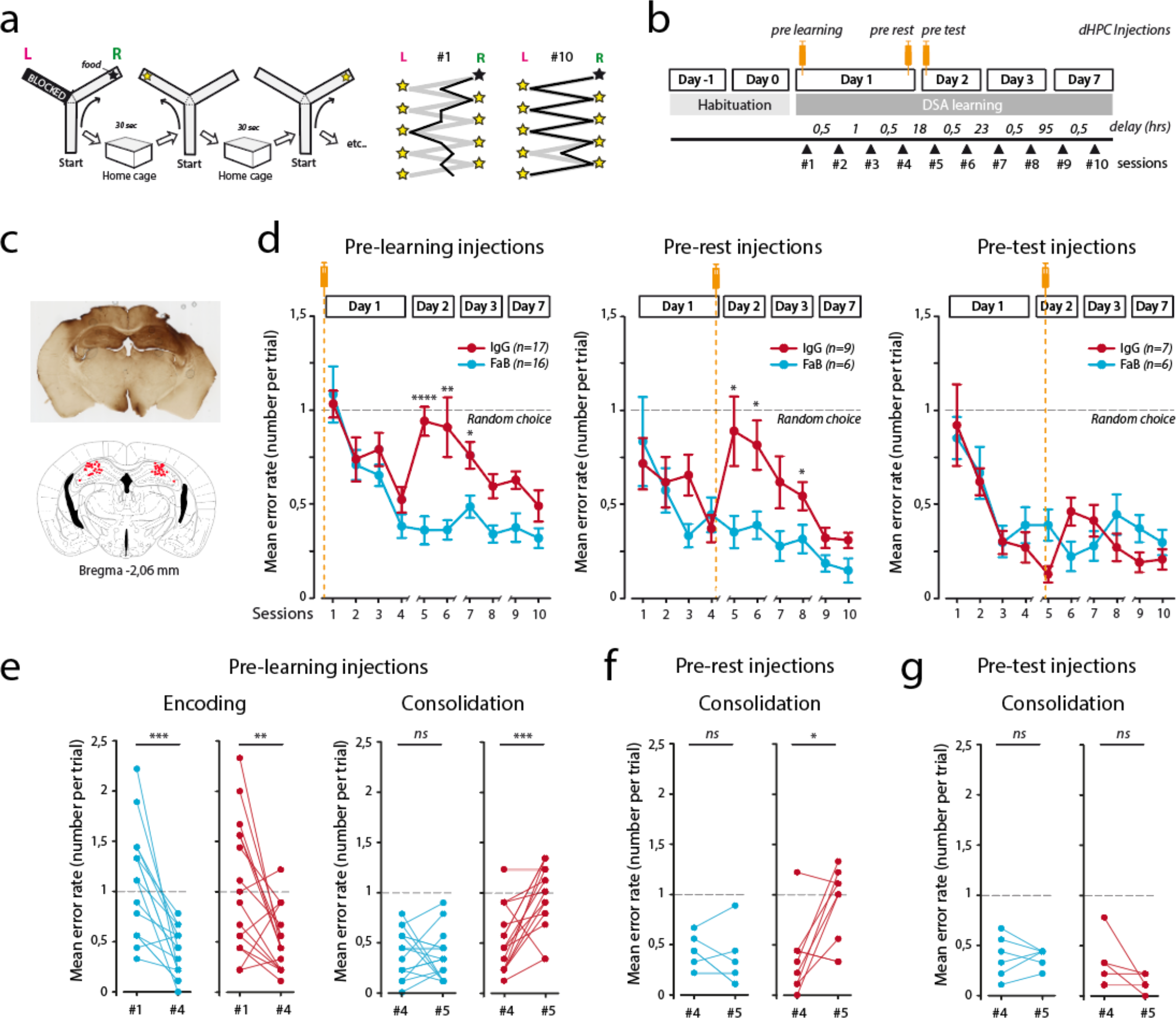
AMPAR surface mobility in the dorsal hippocampus is necessary for consolidation of a delayed spatial alternation rule. **a:** schematic of the DSA (D elayed <u>S</u> patial A lternation) rule that animals have to acquire. In each session, after a first forced choice, nine food-rewarded positions are set, alternatively in the right or left ending arm. Each session has 10 trials, each associated to a given reward position . Up to five error runs are permitted per trial before the animal is forced to enter in the rewarded arm. In between runs, the animal is positioned in its home cage for 30 seconds. After 10 training sessions (session#1 to session#10) allocated within a week, animals are alternating almost perfectly (right panels). **b**: Intracerebral bilateral injections in the dorsal hippocampus were performed at three different time points of the DSA training: pre learning (before session#1), pre-rest (after session#4) and pre- test (before session#5) (syringes). **c**: histological profile of the injections. Two cannulas were implanted above the dHPC, and anti GluA2-IgGs or control compounds injected (top). In order to cover a large portion of the dHPC, multiple injection points were used in the ventral-dorsal axis (middle). Entry points of the injection cannulas were mostly above the CA1 area (bottom). **d**: Behavioural results obtained in the various cohorts expressed as mean error rates. Injections of anti GluA2 bivalent IgGs (red) or monovalent Fabs (blue) were performed before session#1 (pre- learning, Left), after session#4 (pre-rest, middle) or before session#5 (pre-test, Right). *: p<0.05, **: p<0.01, ***: p<0.001. N numbers represent the number of injected animals. **e**: Single animal data are shown for crucial behavioural steps. Memory encoding (left) is supposed to be achieved within the first day (after 4 sessions of 10 trials each) as the error rate is minimal at session session#4 (Figure 1d). Error rates between session#4 and session#5 are similar in FaB- injected control animals, suggesting optimal memory consolidation. In contrast, Error rate returned to chance level in IgG-injected mice, denoting lack of rule consolidation. *ns*: not significant, **: p<0.01, ***: p<0.001. **f-g**: Same presentation as in **e**. Results for consolidation - session#4 VS session#5 – in pre-rest ( **f**) and pre-test (**g**) injected cohorts are shown. *ns*: not significant, *: p<0.05, **: p<0.01. ***: p<0.001.

An important question to address is the origin of the lack of performance observed in day 2 in pre-learning and pre-rest injected animals. Indeed, an increase in the number or errors can have various origins such as disorientation, disengagement, bad animal state, up to the complete forgetting of the alternation rule. To further dissociate between these options, we thoroughly analysed animal behaviour along the DSA rule acquisition ( **Figure 2**). As other groups using T and/or Y mazes to test for mice cognitive abilities ^16, 17^ , we noticed that runs can be separated in two groups: those in which the animals were running in the maze with almost constant speed and those in which hesitation can be observed at the crossing point, with significant changes in head orientation and speed, called <u>V</u>icarious <u>T</u>rial and <u>E</u>rror runs or VTE runs ( **Figure 2a**) that are predictive for good choice accuracy, as testified by the difference in the error rates in VTE and non VTE runs ( **Figure 2a_right_**). Interestingly, probably because no rule was clear initially, animals exhibited more no-VTE runs at the beginning of the learning day (**Figure 2b**).

**Figure 2:**
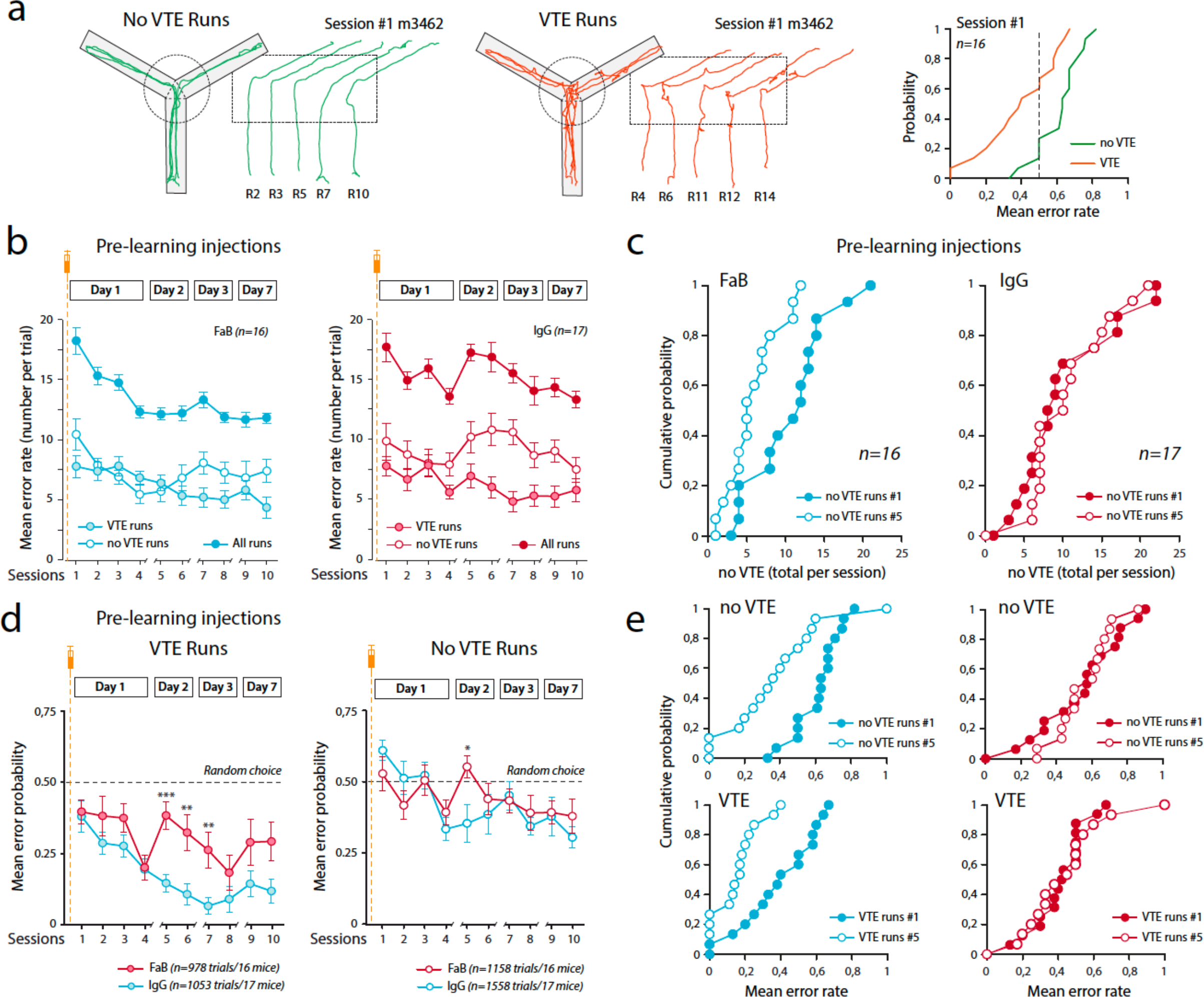
Immobilization of AMPAR in the dorsal hippocampus led to complete forgetting of the acquired DSA rule. **a:** DSA runs can be separated in two groups according to animal behaviour (hesitation) in the middle of the maze. As defined by ^17^, Vicarious Trial and Errors behaviour indicate the cognitive engagement of the mouse in the task. **b**: For pre-leaning injections cohorts, number of VTE and no VTE runs was analysed along the DSA sessions. Note that upon IgG injections, the number of no VTE runs at session#5 was similar as in session#1. **c**: Cumulative single animal data for no VTE run numbers in session#1 and session#5. Note that curves are superimposable following pre-learning IgG injections. **d**: Choice acuteness during VTE and no VTE runs was analysed along the DSA sessions in the pre-learning injection cohorts. The lack of animal performance at session#5 (see Figure 1d) is associated with a strong decrease in VTE runs accuracy that returned to its initial value in session#1. A similar effect is observed for no VTE, with a lesser extent. *: p<0.05, **: p<0.01. ***: p<0.001. **e**: Same presentation as in **c** for choice accuracy at VTE and no VTE run numbers in session session#1 and session#5. Note that curves are superimposable following pre-learning IgG injections.

Next, to get insights in the origin of the performance loss of pre-learning treated animals, we examined the occurrence and choice quality of VTE and no-VTE runs in session#5 ( **Figure 2b-e**): Surprisingly, the lack of performance of pre-learning IgG-injected animals was associated with reinstatement of initial values, suggesting an apparent DSA rule amnesia ( **Figure 2c-e**): **i)** in IgG-treated mice, the occurrence of no-VTE runs in session#5 was similar to its initial session#1 value (**Figure 2b,c**), as if the animal had to re-establish a rule, rather than being unable to apply the beforehand encoded one; **ii)** the error rate of VTE runs in session#5 also increased, returning to the level observed in session#1 ( **Figure 2d**), pointing that when wanting to apply the rule, animals performance was as when encountering it initially ( **Figure 2e_bottom_**) and **iii)** Whereas with training the no-VTE runs performance improved and progressively diverge from random choices, they also returned back to initial values in session#5 of IgG-inject e**F**d**ig**m**ur**i**e**c e**2e**(**_top_**). Importantly, in day1, all these parameters evolved similarly between IgG- and control-injected animals, suggesting that DSA rule encoding process is ongoing normally in absence of AMPARM ( **Figure 2b-e** and **Supplementary Figure 1**). Thus, we propose that a hippocampal AMPARM-dependent mechanism is at play during post-training resting periods that support memory consolidation, and that dHPC AMPAR immobilization leads to a total forgetting of the acquired DSA rule rather than to an incorrectly encoded or played DSA rule.

### AMPAR immobilization in the dorsal hippocampus impairs ripple physiology during slow wave <u>sleep.</u>

Hippocampal ripples are fast oscillations that are developing during slow wave sleep (SWS) phase and that are considered as offline replays of immediately preceding experiences ^6, 8, 18^. They are generated in CA2/CA3 regions of the hippocampus, and propagate in CA1 before broadcasting to cortical regions ^9, 19^. Interestingly, their interplay with immediately preceding synaptic tagging is unknow, even if specific downscaling and NMDAR-dependent synapse refinement have been reported in *in situ* preparations^11^.

Thus, we wanted to examine the impact of IgG treatment and DSA learning on dHPC ripples ( **Figure 3**). To achieve that, animals were implanted bilaterally with wire bundles medially to injection cannula ( **Supplemental Figure 2a**). dHPC Local Field Potentials (LFPs) were recorded for 3 hours immediately following Y maze habituation (“habituation” in Day-1 or D-1) or 4 first DSA sessions (“DSA” in Day1 or D1, **Figure 3**). At first, we separated awake and resting/sleeping state in the home cage using animal tracking (mobility, **Figure 3 _top_** and **Supplemental Figure 2c**). Then, Slow Wave Sleep (SWS) and Rapid Eye Movement (REM) sleeping phases were separated using a Theta/Delta ratio defined on hippocampal LFP spectra ( **Figure 3_middle_** and **Supplemental Figure 2c**), REM periods hosting robust theta oscillations absent in SWS periods, that are characterized by pronounced Delta waves (for a typical example see **Supplemental Figure 2c**). Importantly, as expected, SWS periods correlate nicely with the occurrence of hippocampal ripples (see methods, **Figure 3 _bottom_** and **Supplemental Figure 2c**).

**Figure 3:**
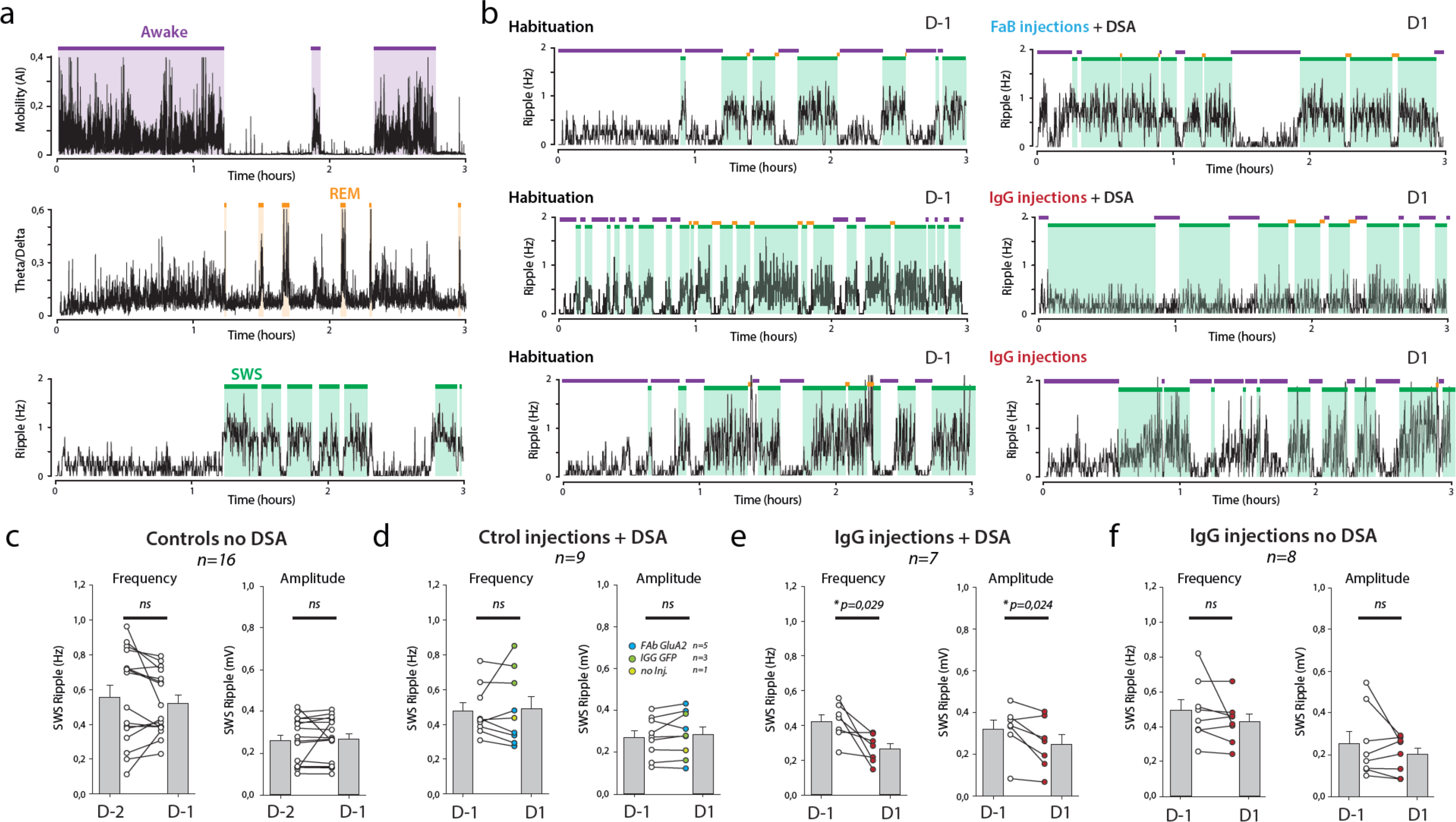
Immobilization of AMPAR in the dorsal hippocampus led to learning-dependent impairment of ripple activities. **a:** During resting periods in the home cage, animal tracking and dHPC electrophysiological LFP recordings were performed, to temporally define three animal states: awake state is defined by animal mobility, whereas resting states were separated in REM (R apid <u>E</u> ye <u>M</u> ovement) and SWS (<u>S</u>low <u>W</u> ave <u>S</u>leep) pahses, according to the Delta/Theta ratio of dHPC LFPs. LFP signals were also filtered at 150-250 Hz to extract ripples, the frequency of which is closely correlated with SWS periods^6^. **b**: Bilateral dHPC LFPs were recorded for three hours resting periods before (habituation, D-1) or after DSA encoding (after session#4, D1). Typical examples of ripple frequency in pre-learning FaB (Top) and IgG injected (middle) animals, or in IgG injected animals that were not subjected to DSA learning (bottom). Note the decrease in SWS-ripple frequency in D1 of the IgG injected DSA-trained animal. **c-f**: Amplitude and frequency of ripples during SWS periods were extracted in D-1 and D1 resting periods in four different groups: **c**: day to day recordings with no injection nor DSA learning. **d**: before and after control drug injections and DSA learning, **e**: before and after IgG injections and DSA learning, **f**: before and after control IgG injections but no DSA learning. *ns*: not significant, *: p<0.05. Number of recorded animals is indicated.

Then we tested if the DSA protocol and AMPAR cross-linking were leading to alterations of ripple frequency and amplitudes ( **Figure 3c-f**). At first, we tested if our recordings were stable over time: indeed, and not surprisingly, neither the amplitude nor the frequency of detected ripples differ between two basal consecutive days (Controls D-2 and D-1 recordings, **Figure 3c**). Some reports described that spatial learning or retrieval was leading to increases in dHPC ripple frequency ^20^.

However, no noticeable changes in recorded ripples were observed in animals submitted to DSA learning, and injected with control constructs or non-injected (see specific mentions in **Figure 3d**). We next tested if the blockade of AMPARM in the dHPC was perturbing ripple physiology in DSA trained animals ( **Figure 3e**) or non-trained mice ( **Figure 3f**). Importantly, in both cases, as for control injections, IgG injections were done several hours before the recorded resting periods, the only difference being that non-trained animals are solely positioned in the maze for a similar time duration, but with no specific rule to learn (see methods). Surprisingly, we detected a significant impact of IgG injections on ripple amplitude and frequency that were significantly decreased, but only when learning was present (compare **Figure 3e** and **3f**). Because ripple inactivation during SWS has been proved to impair spatial memory consolidation ^21^, this decrease in ripple content may explain the lack of consolidation observed in pre-learning IgG injected animals ( **Figure 1d**). Importantly, we did not detect any change in the total time of SWS between groups in D1 within the first three hours of resting period (data not shown). Importantly, this early SWS phase has been shown to elicit ripples that are of particular importance for recent memory consolidation ^18^. Thus, our data support that in the dHPC, DSA-consolidation mobilize AMPARM-dependent plasticity events that support the genesis and strength of hippocampal ripples.

### AMPARM-dependent plasticity at CA3 recurrent support ripple activity *in situ*

Most of what we know about cellular and synaptic contribution to ripple physiology comes from acute *in situ* preparations in which ripple like oscillations are spontaneously generated ^10, 12, 13^. If some aspects of *in vivo* ripples are obviously lacking, such as their cognitive content ^6^, they recapitulated most of the *in vivo* ripple properties and keep some link with *in vivo* experience^116^.

We setup and used *in situ* hippocampal preparations exhibiting spontaneously ripples and combined them with synaptic “tagging” by inducing LTP at CA3 ➔CA1, CA3 ➔CA3 and DG ➔CA3 synaptic contacts with or without presence of AMPAR cross-linkers ( **Figures 4 and 5**). Among the addressed questions is the interplay between AMPARM-dependent plasticity and ripple physiology to get insights on our *in vivo* results. In optimized *in situ* preparations ^13^, ripple-like activities – here called SPW-Rs – can be stably and robustly recorded using field recording pipettes positioned in the CA3 and CA1 regions ( **Figure 4a-e**). To be included, SPW-Rs recordings has to have stable occurrence frequency, showing co-detected CA3 and CA1 events, present a constant delay between CA3 (first) and CA1 (delayed) responses (**Figure 4c _middle_**), and have a good amplitude matching between both signals ( **Figure 4b and 4c _right_**). Some other criteria were eventually respected when present: **i)** the signal polarity in the CA1 region was depending on the recording location: positive in the *stratum pyramidale*, and negative in the *stratum radiatum* , confirming that incoming CA3 activities were generating a significant synaptic field response in CA1 (**Supplementary Figure 3a**), **ii)** both evoked and spontaneous SPW-Rs eventually engaged CA3 unitary activities ( **Supplementary Figure 3b**), **iii)** when tested, stimulations in the CA1*stratum radiatum* that generated SPW-Rs were interfering with spontaneous SPW-Rs, generating a refractory period ( **Supplementary Figure 3c**). Importantly, after a 20 minutes’ period in the recording chamber, all parameters were stable for more than an hour, allowing the combination of SPW-Rs recordings with HFS application and/or pharmacological manipulations (**Figure 4 and 5**).

**Figure 4:**
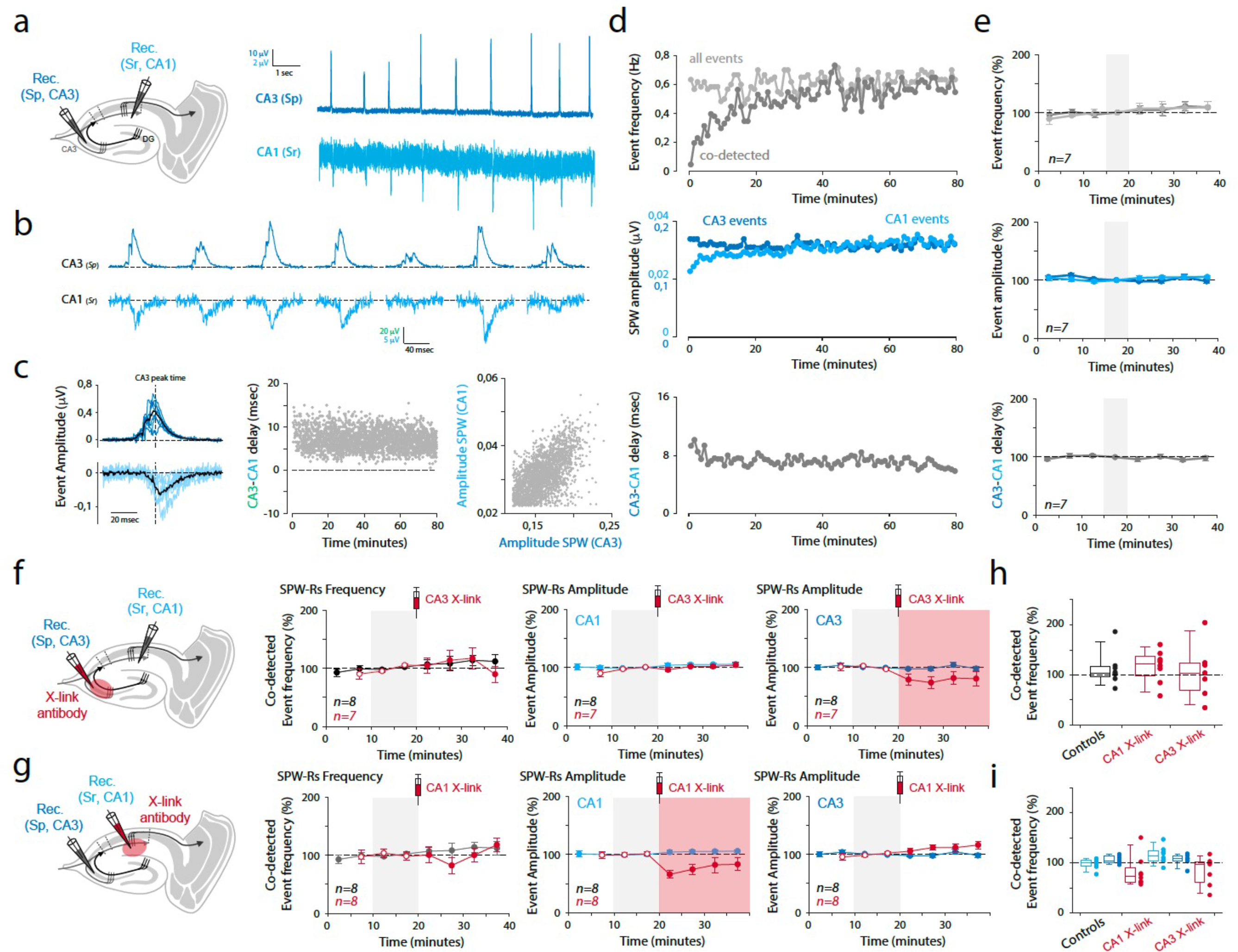
Immobilization of AMPAR in the dorsal hippocampus did not affect spontaneous ripple activities in naive *in situ* preparations. **a:** Spontaneous Sharp waves events (SPW-Rs) are recorded in fresh *in situ* hippocampal preparations (see methods) using extracellular field electrodes**b**. : Examples of recorded events. **c**: Simultaneously recorded events showed a significant delay, and correlated amplitudes denoting their propagation from CA3 to the CA1 area. **d-e**: Single example (d) and averaged (e, n=7 similar experiments) measures showing the stability of SPW-Rs frequency (Top), amplitudes (middle) and delay (bottom) with time. All pharmacological experiments are starting after respecting a 20 minutes period required for SPW-Rs stabilisation. **f-g**: Effect of AMPAR X-linking on spontaneous SPWRs was tested in in situ preparations by pressure injection of anti-GluA2 IgGs. Left: schematic of the experiment. Right: Time course of SPW-Rs frequency (CA3-CA1 co-detected events) and amplitude in CA1 and CA3 regions. The “light red” area indicate that events are recorded in the IgG-injected area. All except red colour indicate time course of the same parameter in control conditions. Open red circles are the IgG preparations before the injections. **h-i**: All single experiments and average values after 20 minutes, in control or after IgG injections. Note that a significant decrease in SPW-R amplitude is observed at the locus of IgG injection (see **supplementary** Figure 5).

**Figure 5:**
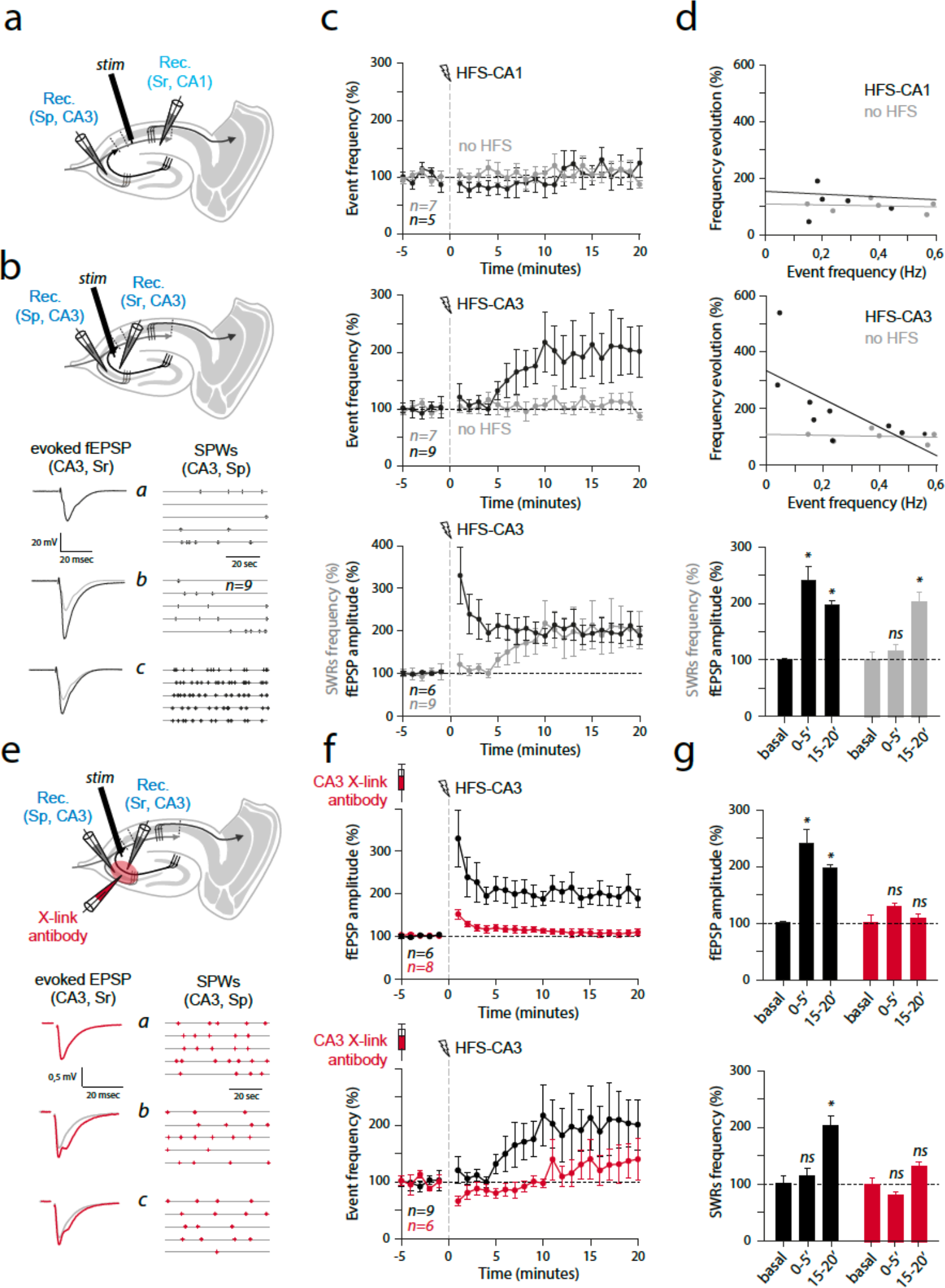
Interplay between plasticity induction, spontaneous SPW-Rs and AMPAR mobility in hippocampus *in situ*. **a:** Spontaneous Sharp Waves Ripples were recorded in hippocampal acute slices. After stabilisation, application of HFS in the *stratum radiatum* of CA1 (CA1-sr) was applied to induce LTP a➔t CCAA31 synapses ( **see Supplementary** Figure 4). No major impact of CA1-HFS onto SPW-Rs frequency or amplitude was observed. Grey zone: time period for baseline calculation; Yellow zone: time after HFS application. **b:** Top: same presentation as in **a**, for HFS application in CA3-sr. Note the increase in SPW-Rs frequency and amplitude after HFS application. Bottom: Left: Typical example of the effect of CA3 HFS on evoked CA3 responses, and frequency of spontaneous SPW-Rs. A significant delay exists between synaptic potentiation (evoked _f_EPSCs, middle; SPW-Rs amplitude, right) and SPW-Rs frequency increase that can be better appreciate in the time courses expressed by minutes. **c**: Results presented in a and b are summarized. **d**: Top: the effect of CA3 HFS (black dots) depends on the initial SPW-Rs frequency. SPW-Rs frequency evolution with time did not depend on initial frequency (grey dots). Bottom: A tendency for a co-evolution exist between CA3-HFS effects on SPW-Rs frequency and amplitude. **e**: Same presentation as in **b**. Before electrophysiological recordings and CA3-HFS applications, IgG injections were performed in the recorded CA3 region (CA3 X-link antibody). Note that all effect triggered by CA3-HFS in control conditions are absent in presence of the IgG. Number of recordings is indicated. **f-g**: Same presentation as in **c** and **d**. The experiments run in presence of AMPAR X-linkers were added. *ns*: not significant, *: p<0.05.

We previously showed that AMPAR immobilisation at neuronal surface in the CA1 region impaired LTP expression at CA3 ➔CA1 synapses^14^. We first wanted to reproduce and extend this finding to other synapses eliciting post-synaptic LTP expression. Interestingly, in CA3 region, pyramidal neurons receive two major excitatory afferent that are intrinsically different: Mossy fibres originating in the dentate gyrus generate “detonating” synapses expressing a huge rate-dependent facilitation that can be prolongated by a sustained potentiation of presynaptic origin (presynaptic release probability increase^22^). In contrast, recurrent synapses emitted by distant or neighbouring CA3 pyramidal cells are classical Hebbian synapses, expressing post-synaptic LT^2^P^2^. We thus tested the impact of AMPAR cross-linking on LTP in our *in-situ* “SPW-Rs” preparation ( **Supplementary Figure 4**). Not surprisingly, we observed that AMPAR X-linking led to an absence of LTP at synapses postsynaptic to CA3 axons (CA3➔CA1 and CA3 ➔CA3) but not at DG ➔CA3 projections, that were solely affected by PKA blockade using Rp-cAMP preincubations ( **Supplementary Figure 4**).

In the absence of learning-associated synaptic tagging, AMPAR immobilization did not lead to changes in ripple frequency or amplitude ( **Figure 3f**). Acute slice preparations from naive mice are often used as models to study molecular and cellular mechanisms of LTP induction and expression at hippocampal synapses ^23^. I t i s t h e n o f t e n c o n s i d e r e d t h a t i n n a i v e m i c e , n o s i g n i f i c a n t tagging – endogenously triggered LTP - is present. We thus tested the effect of AMPAR cross-linking in SPW-Rs containing naive preparations by locally infusing anti-GluA2 IgGs in CA1 or CA3 *stratum radiata* (**Figure 4f-i**). Importantly, the efficacy of this injection procedure on LTP expression was previously validated in CA1^14^, and reiterated here in CA1 and CA3 regions **S**(**upplementary Figure 4**). As compared to basal conditions, these injections had no effect on SPW-Rs frequency or amplitude (**Figure 4f-i**), the local effect of IgG injection on amplitude being attributable to the one/two pipette(s) procedure ( **Supplementary Figure 5**). Thus, in great coherence with the absence of AMPAR immobilization on basal synaptic transmission ^14^, and *in vivo* results obtained in naive mice, our *in-situ* result suggest that basal SPW-Rs did not depend on AMPARM in absence of specific synaptic tagging.

Next, we wanted to test for the effects of synaptic tagging – here generated by HFS applications enabling LTP induction ( **Supplementary Figure 4**) – on SPW-Rs frequency ( **Figure 5**). Interestingly, CA1 HFS stimulations did not impact SPW-Rs frequency or amplitude ( **Figure 5a, c, d**), indicating that synaptic strength at CA3 ➔CA1 synapses may not be a prominent SPW-Rs frequency determinant. In contrast, the same procedure applied at CA3 ➔CA3 recurrent synapses led to a strong increase in SPW-Rs frequency ( **Figure 5b, c**), prominent in case of low basal SPW-Rs frequency ( **Figure 5d**).

Furthermore, the effect of HFS on synaptic strength and SPW-Rs frequency seems to be temporally disconnected, the increase in evoked EPSP amplitude being detectable as early as in the 0-5 post-tetanic period, whereas the effect on SPW-Rs frequency was not yet present ( **Figure 5d_bottom_**). This suggest that the reinforcement of CA3 ➔CA3 recurrent synapses progressively increase CA3 region excitability, more prone to generate ripples.

Finally, we tested the impact of AMPAR immobilization on SPW-Rs modulations by CA3-HFS to test if they would be independent. We applied HFS-CA3 stimulations in SPW-Rs expressing slices in which local CA3 infusions of anti GluA2 IgGs were performed ( **Figure 5e-g**). Under AMPAR immobilization, as found for LTP at CA3 ➔CA3 synapses, HFS-associated effect onto SPW-Rs frequency was absent (**Figure 5e-g**), suggesting that their physiology depend on AMPARM-dependent CA3 recurrent synaptic strength. Importantly, a contribution of synaptic potentiation at DG ➔CA3 synaptic inputs is unlikely, as we systematically tested for 1Hz frequency facilitation in our evoked synaptic responses (Data not shown), and also because they arbour a form of plasticity that is insensitive to AMPAR cross-linking ( **Supplementary Figure 4**). Thus, we would like to propose that AMPARM-dependent LTP at CA3 recurrent synapses positively controls ripple activity *in situ*. Together with *in vivo* data, it suggests that CA3 ➔CA3 synaptic tagging is triggered during DSA learning, allowing ripple-mediated consolidation to occur during consecutive sleep phases.

### AMPAR mobility in CA3 area is necessary for memory consolidation

Based on our *in situ* data, we ambitioned to restrict AMPAR cross-linking to the CA3 area to evaluate if impairing plasticity only in this region would be sufficient in impairing memory consolidation and ripple physiology. However, specific antibody-based AMPAR cross-linker strategy lacks of spatial and temporal resolution: Indeed, to maximize its efficiency, multiple *in vivo* injections were performed on a dorso-ventral axis to cover most of the dHPC ^14^. In addition, secondary, unwanted effects of anti-GluA2 antibodies on AMPAR composition ^24^ may be present that can lead to misinterpretation of the data ( **see discussion**). Thus, we used a recently developed approach to cross-link endogenous GluA2-containing AMPAR using biotin/streptavidin complexes ^15^ (**Figure 6a**). In Knock-in mice expressing AP-tagged GluA2 subunits, the presence of an exogenous enzyme – BiRA ^ER^, brought by viral infections – allow the biotinylation of GluA2-containing AMPAR, that can be cross-linked in the presence of tetravalent streptavidin added in the extracellular space ( **Figure 6a**). This cross-linking approach that has been validated *in vitro* and *in vivo* ^15^ and among other advantages, will help in getting improved spatial resolution, as combining viral expression and drug delivery through intracerebral cannula (**Figure 6b**). We tested our capacity in targeting CA3 area by infusing NA- texasRed (red-tagged tetravalent neutravidin through cannula implanted above the CA3 regions of BIRA-expressing mice ( **Figure 6c**). Indeed, red labelling was almost completely restricted to the CA3 region, in a subpart of the green expressing region ( **Figure 6c**). Then, we tested the capacity of mice in retaining DSA rule in various control and CA3 cross-linking conditions. Because the time course of NA action is not yet known, we privileged pre-rest injections ( **Figure 6b**), and GluA2 KI animals being slow in establishing alternating behaviour, we mixed sessions #1-2 and #5-6 to get more robust behavioural outcomes. When compared to session#1-2 – a time point at which no learning is achieved – the error rate of control animals was significantly l o **F**w**ig**e**u**r**re**i**6**n**d** s) essions #5-6 ( indicating that encoding and consolidation have been successfully achieved. In contrast, error rates at these two time points remain close to random values in X-linking conditions ( **Figure 6d**) a phenotype that is again accompany by an apparent forgetting of the DSA rule, as the accuracy of VTE runs, that improved in control mice, did not show any evolution in the X-linking conditions ( **Figure 6e**).

**Figure 6:**
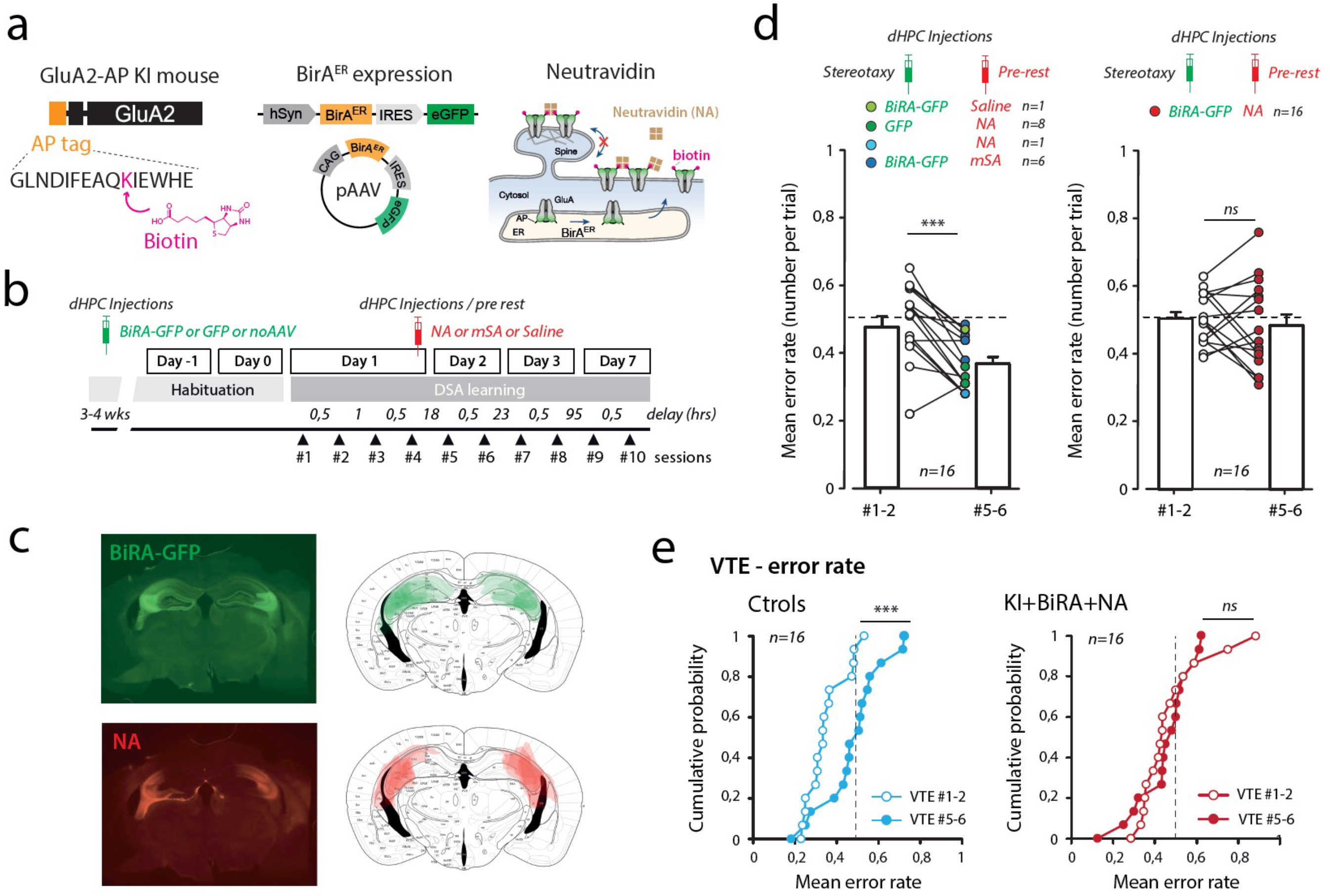
An alternative AMPAR X-linking strategy allowing a better targeting to the CA3 area also induced complete forgetting of DSA rule. **a:** We recently developed a new strategy for AMPAR X-linking. Knock-in mice expressing endogenous AP-tagged GluA2 AMPAR subunits can be biotinylated in presence of BiRA ^ER^, and once exported to the cell surface can be immobilized in presence of external neutravidin (NA, cross-linking condition). **b**: similar *in vivo* pharmacological experiments as in Figure 1d were performed, combining early stereotaxic dHPC injections of AAV-BiRA-GFP or AAV-GFP, and pre-rest injections of saline, mSA or NA. **c**: histological controls for the mSA and NA staining on top of the AAV-GFP expression. The combination of both injections better restrict AMPAR immobilization to the CA3 area. **d**: Mean error rates were compared between session session#1 and session session#5 to evaluate the retention of the DSA rule upon various pharmacological treatments (as indicated by colour coding). **e**: As in Figure 2, we reported the number of no VTE runs that were observed in sessions session#1 and session#5. The amount of no VTE runs was similar between sessions session#1 and session#5 in the KI/BiRA/NA condition (right) pointing for memory forgetting, whereas different in the control group (Left). **f**: The error rate in VTE runs was analysed and reported in sessions session#1 (Left) and 5 (Right) for control and cross-link groups. Note that they were identical in Session session#1, but diverge significantly in session session#5, suggesting memory impairment in absence of AMPARM. *n s*: not significant. ***: p<0.001.

To confirm that the lack of consolidation is associated with impairment of ripples physiology, we combined this novel cross-linking method with dHPC recordings using the same recording methodology as for IgG experiments ( **Figure 7**). Ripples occurring during SWS were extracted and counted in habituation (D-1) and after DSA (D1) sessions ( **Figure 7**). This last recording starting immediately after drug delivery, it was important to control for unspecific effects of drug actions: importantly, NA application on GFP-only, and saline or mSA delivery on BiRA expressing dHPCs were not leading to changes in ripple frequency, mirroring the lack of effect on DSA consolidation ( **Figure 6**). However, NA delivery in BiRA expressing CA3 areas were associated with a pronounced decrease in ripple frequency, remembering the effect observed upon pre-learning and pre-rest IgG injections (**Figure 3**). Thus, by two different approaches, we demonstrated that AMPARM in the CA3 region is necessary for memory consolidation and support ripple physiology during slow wave sleep.

**Figure 7:**
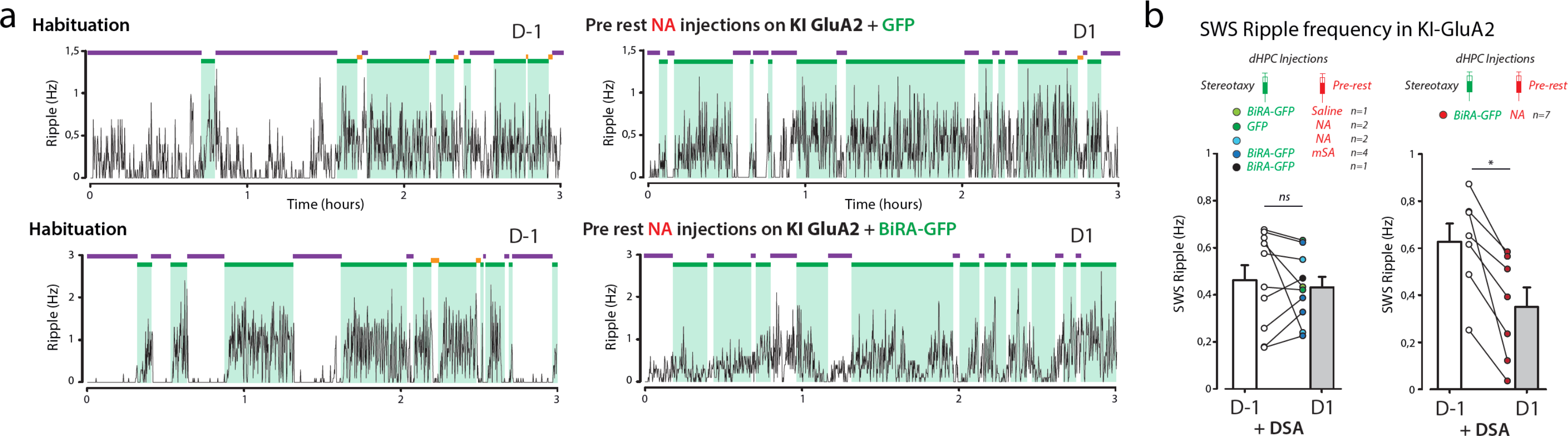
An alternative AMPAR X-linking strategy allowing a better targeting to the CA3 area affected SWS ripple activity. **a**: Same presentation as in Figure 3b. Bilateral dHPC LFPs were recorded for three hours resting periods before (habituation, D-1) or after DSA encoding (after sessions session#4, D1). Typical examples of ripple frequency in control (Top and bottom) and X-linking (middle) conditions. Note the decrease in SWS-ripple frequency in D1 of the KI-BiRA-NA DSA-trained animal. **b**: Same presentation as in Figure 3c. SWS ripple frequency was analyzed and reported for single experiments (dots) and averaged (bars). Conditions and animal numbers are indicated. *ns*: not significant, *: p<0.05.

## Discussion

In order to understand the link between synaptic plasticity and learning and memory it is essential to analyse separately the various phases of memory encoding, consolidation and retrieval, and to use specific tools that disturb plasticity without affecting basal synaptic transmission. With this in mind, we develop molecular strategies that can be used *in vivo* to address these issues. We tested two different methods to impair postsynaptic long-term potentiation in the dorsal hippocampus, and uncover that during the process of acquiring a spatially guided rule to get food rewards, AMPAR mobility was necessary during the consolidation phase and was an important physiological mechanism that support ripple activity consecutive to new rule encoding. From our *in-situ* experiments, an interplay between AMPARM, LTP at CA3 ➔CA3 recurrent synapses and ripple physiology emerged, suggesting a model ( **Figure 8**) in which rule encoding, possibly through synaptic tagging, condition/organize subsequent ripple activity that will develop during rest to consolidate memory. This would require the occurrence of AMPARM-dependent LTP at CA3 recurrent synapses, as the immobilization of AMPAR in*in*C A*vi*3*vo* during consolidation leads to a learning-dependent loss of ripple activity and complete forgetting of the acquired rule. Thus, we bring a new mechanism by which synaptic plasticity contribute to learning and memory.

**Figure 8:**
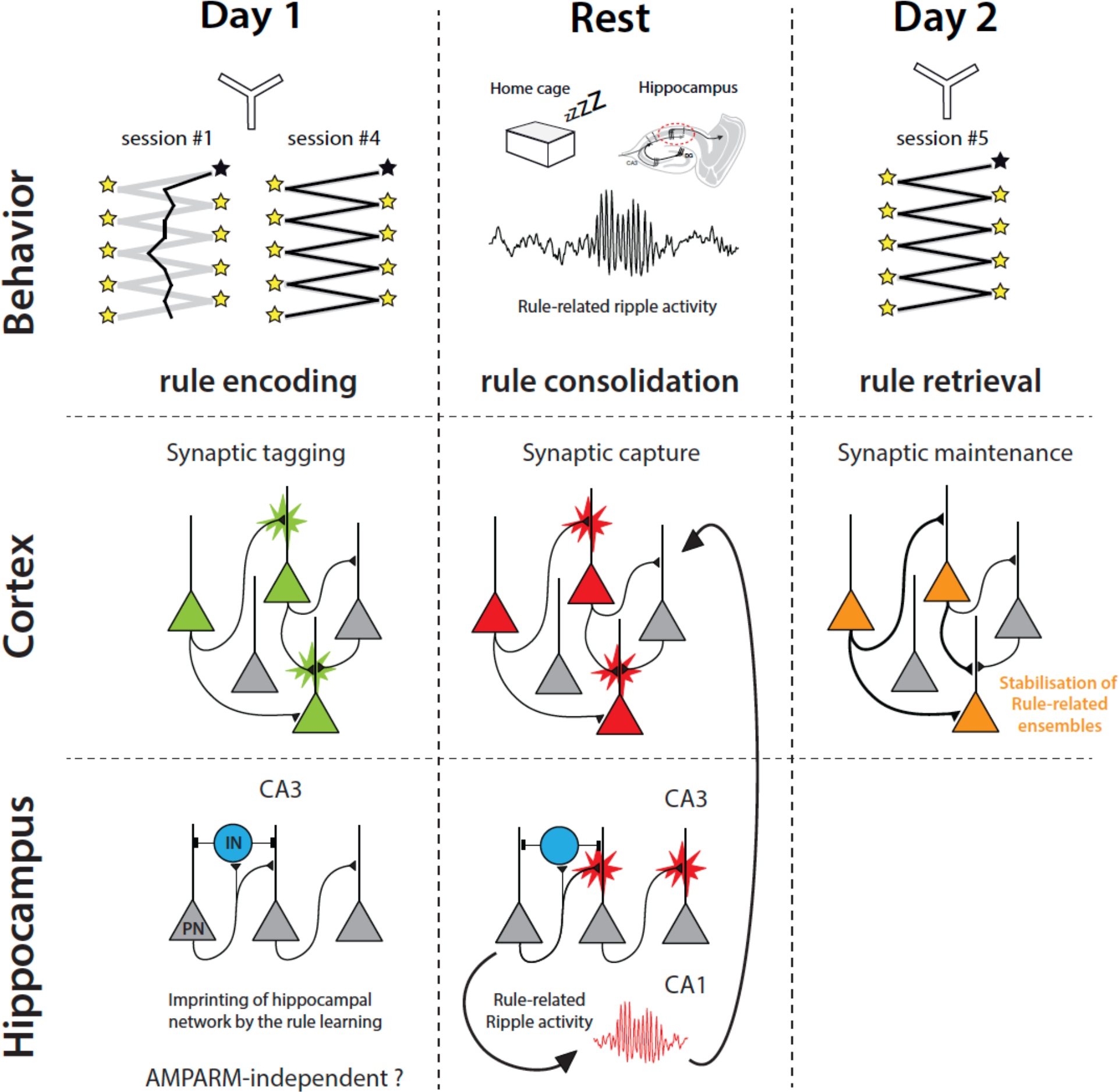
Working model. Model proposed for the action of AMPAR X-linking onto DSA memory consolidation. DSA rule encoding lead to synaptic tagging in the cortical areas, including the mPFC. Hippocampal remapping that is possibly occurring would remain insensitive to GluA2-dependent AMPAR immobilization (EX-IN CA1 ➔CA1 synapses or DG ➔CA3 synapses). During SWS, ripples necessary for synaptic capture in the cortical areas (through replays-dependent reactivation of neuronal ensembles) would be impaired as plasticity at CA3 ➔CA3 recurrent is impaired.

### Controls for AMPAR X-linking strategies

Some antibodies against GluA2 subunits have been reported to modify AMPAR composition within several hours ^24^, a time windows that may correspond to the consolidation phase of the DSA rule in case of pre-learning injections. Thus, one could attribute the observed effects onto consolidation to important changes of hippocampal connectivity/activity due to the replacement of GluA2-containing heteromers by GluA1 homomers. However, we observed a very similar effect on animal performance when intracerebral IgG injections were performed before or after DSA rule encoding ( **Figure 1d-f**), a time at which the IgG-dependent changes in AMPAR composition might not have yet occurred. The same reasoning applies to the ripple recordings in **Figure 3**: The application of pre-learning IgG in absence of DSA learning did not impact the network capacity in generating ripples. In fact, the impact of both strategies blocking AMPARM appears to be specific on the actual cross-link capacity (see the various control conditions for both antibody- and neutravidin-based strategies) at the time of offline rule consolidation.

A methodological issue regarding *in vivo* strategies and ripple detection is that local delivery of drugs affects the electrophysiological recordings due to volume injection, an issue that was identified in our *in-situ* e x p e r i m e n**su**t**p**s**pl**(**ementary Figure 5**). If occurring *in vivo* , a global decrease in LFP amplitude caused by tissue movements would have potentially affected our capacity to detect ripples after injection. At first, we tried to minimize this effect by decreasing the injection speed (see methods) and temporally separate the injection from the recordings (pre-learning injections). We also designed a number of control conditions that would account for this effect: we injected animals with either saline, various monovalent or divalent antibodies, but also with IgG without submitting the animals to DSA learning ( **Figure 3**). In all of these conditions, no changes in ripples properties were observed.

### Comparison between *in vivo* and *in situ* recordings

Another issue concerns the comparison of SWRs recorded *in vivo* and *in vitro* ^6^. Indeed, in our *in vivo* recordings, we essentially characterized changes in frequency and amplitude of oscillations recorded in the CA1 region after a 150-250 Hz band pass filtering ( **Figures 3** and **7**), whereas focusing on the occurrence frequency of CA3/CA1 sharp waves *in situ* (SPW-Rs, **Figures 4** and **5**). Sharp-waves are proposed to reflect the dendritic depolarization evoked by the synchronous activity of subgroups of excitatory afferents from the CA3 region, whereas the ripple oscillations are thought to be generated in the CA1 region, in response to the sharp-wave-associated excitatory inputs ^12^. In the retained *in-situ* recordings, the vast majority of SPW-Rs are co-detected in CA3 and CA1 regions with a significant and stable delay ( **Figure 4**). In CA1, a 150-250 Hz band pass filtering of SPW-Rs eventually give rise to a ripple-like oscillation (**supplementary Figure 3a**), but that appears to be highly unstable. Another issue is that in some CA3 *stratum pyramidale* r e c o r d i n g s , 1 5 0 - 2 5 0 H z r i p p l e s w e r e c o n t a m i n a t e (**supplementary Figure 3c**). Thus, the use of 150-250 Hz ripples *in situ* appears to be less reliable than the associated co-detecte d*In*w*vi*a*vo*v e, st .he presence of dendritic responses – waves – is present in some but not all recordings, as the wires implanted are separated by 200 μm in the dorso-ventral axis, thus radially to the CA1 region ( **supplementary Figure 2a**). However, because waves and ripples are reflecting the same intrinsic network events, we believe that effects observed on *in vivo* ripples and *in situ* sharp waves frequencies can be compared, especially the fact that they share the same lack of sensitivity to AMPAR X-linking in basal condition, whereas being impaired by the same treatment after DSA learning ( *in vivo,* **Figures 3** and **7**) or LTP induction (*in situ,* **Figure 5**).

### Synaptic plasticity in memory phases

Surprisingly to us, the effect of AMPAR X-linking on ripple activity is present only if the DSA rule is encoded ( **Figure 3**). This suggests that ripple activity is different if salient cognitive events have to be consolidated. One intriguing possibility would be that the learning process directly or indirectly “conditions” or “imprints” the hippocampal network, especially the CA3➔CA3 recurrent synapses, to drive DSA-related ripples to be efficiently generated offline ( **Figure 8**). A likely mechanism by which this could happen is the occurrence of synaptic “tagging” –activity-dependent synaptic plasticity events – during online rule acquisition, that will have to be “captured” during the offline consolidation phase of memory ^4, 5^. It has yet been suggested that *in vivo* synaptic modifications would protect these synaptic contacts from ripple-mediated synaptic downscaling during sleep, allowing cognitive map refinement ^11^. Several aspects of our findings need to be discussed regarding this conceptual line, especially the fact that memory encoding seems to occur even if synaptic plasticity is blocked during rule encoding.

Synaptic tagging is thought to rely on cellular and molecular mechanisms associated with synaptic plasticity induction and expression, that includes AMPAR trafficking at the plasma membrane ^1^.

However, we yet reported that AMPAR cross-linking was not impacting LTP induction ^14^ as NMDA receptors are kept functional. Thus, coincident neuronal activations may have activated transduction cascades necessary for synaptic tagging even if no LTP was expressed.

It remains quite surprizing that we did not observe an impact of hippocampal AMPAR cross-linking on DSA rule encoding ( **Figure 1d _left_**). It would suggest that synaptic tagging associated with rule acquisition does not depend on activity-dependent changes in synaptic strength in the hippocampus, a structure that is assumed to bind all components of experience into one unique episodic memory^25^.

A first hypothesis would be that rule encoding is independent on AMPARM-dependent Hebbian plasticity. Some report yet described that spatial map reorganisation upon rule acquisition can be independent from NMDA-dependent plasticity: Dupret and colleagues ^26^ injected rats with an NMDAR antagonist in order to interfere with their spatial memory. Learning performance was unaffected, but the animals failed to remember the newly-learnt locations, suggesting that the newly-acquired representations of goal locations, that when replayed during sleep predicted memory performance^18^, did not stabilize.

A second explanation would be that our cross-linking strategy did not perturb some hippocampal synaptic contacts crucial for rule encoding. Our two strategies target GluA2-containing AMPAR. Therefore, a number of excitatory synapses may have escaped from the effect of cross-linking, such as excitatory inputs onto interneurons that can be GluA2 independent ^27^. It is of note because some pyramidal cell-interneuron coupling changes have been reported during spatial rule encoding ^28^.

Thus, to have a complete and comprehensive view of AMPARM impact onto learning-dependent ripple physiology, further experiments using cross-linking strategies targeting GluA1-containing AMPAR, or specific cell types for example using conditional expression of the BiRA under IN or PN promoters will be necessary. Along the same line, we observed that DG ➔CA3 LTP was preserved in presence of anti GluA2 IgG, that would possibly support rule encoding and relay broad synaptic tagging within the CA3 region. Further *in vivo* experiments using specific blockers of DG ➔CA3 plasticity, such as Rp-cAMP ( **supplementary Figure 4**) would allow deciphering the role of these particular connections in the encoding of the DSA rule in the hippocampus.

Another possible mechanism for this unexpected disconnection between learning ability and dHPC LTP blockade could reside in the resilience of the system. A recent study using another strategy to block post synaptic LTP showed that even if CA1 plasticity was largely absent, and some of the learning-induced cellular rearrangements were lost, animals were still able to perform correctly ^2^. Alternatively, the rule can first be encoded in another brain region than the dHPC, such as being hosted in the mPFC. For example, Peyrache and colleagues showed that in very similar conditions of a new rule learning in a Y maze, ripple activity was directed towards the reactivation of rule-related neuronal ensembles in the mPFC ^8^, opening the possibility that synaptic tagging have been generated there. Indeed, replay of firing patterns in hippocampal neuron ensembles during sleep is thought to cause the gradual formation of stable representations in extra-hippocampal networks by enhancing connectivity between their elements ^25^. Thus, we can anticipate that pre-learning AMPAR X-linking experiments in the mPFC, by interfering with LTP-dependent synaptic tagging during encoding, would impair animal performance. Similar results would be obtained at pre-rest injections, by blocking ripple-mediated generation of DSA rule representations ^8^. In our hands, complete pre-learning inactivation of dHPC or mPFC through bilateral pre-learning injections of muscimol was leading to a prominent – above chance level - number of errors linked to stereotyped choices (data not shown). This suggests that both structures are necessary to build the cognitive representation of the DSA rule. Importantly, this phenotype was not observed upon IgG-based pre-learning AMPAR cross-linking in dHPC, supporting that no such major effect on dHPC physiology was associated to the procedure.

### Synaptic plasticity and memory maintenance

One interesting observation emerging from our data is the fact that in absence of correct consolidation, the DSA rule is apparently completely forgotten, as if the animal would be completely naive to the task ( **Figure 2**). So far, experiments impacting ripples activity during sleep impaired the performance of the animal on the following days, but were not reported to lead to complete resetting, but more slowing down the behavioural performance ^28^. An intriguing possibility would be that in absence of synaptic capture, synaptic tagging would fade away, bringing the neuronal network in the exact same “naive” state as the previous day, as yet suggested by Norimoto and ^1^c^1^.oInltleereasgtinugelysenough, this would open a time window during which newly encoded memory would be accessible to erasure if one can specifically act onto ripple physiology.

## Conclusion

This study is bringing a first piece of evidence that consolidation of recently acquired memory depends on AMPARM in the hippocampus. Our results points to the importance of CA3 region in this process. Our results are embedded in the more global framework of the tagging and capture synaptic hypothesis that is now more and more discussed in term of encoding / consolidation of memory and awake / sleeping state of the animals. The importance of the cortico / hippocampal dialog in this process is of fundamental interest, and the deciphering of its intimate mechanisms will certainly profit from the development and use of *in vivo* applicable molecular strategies interfering with plasticity*in vivo* with good temporal and spatial control.

## Acknowledgment

We thank T-A. Vernoy, G. Dabee, E. Normand, P. Costet, M. Dehors, C. Martin, for support with animal husbandry and in vivo experiments; S. Marais for imaging and analysis support. Funding: This work was supported by European Research Council (ERC) grants to D.C. (grants ADOS 339541 and Dyn-Syn-Mem 787340), a Fondation Recherche Médicale (FRM) grant to Y.H. (grant DEQ20180339189 AMPA-MO-CO), an Agence Nationale de la Recherche (ANR) grant to Y.H. (grant OptoXL ANR-16-CE16-0026), This work was supported by the Bordeaux Neurocampus core facilities (LabEx BRAIN; grant ANR-10- LABX-43), including the In Vivo Experimental (PIV-EXPE) facilities of the IINS; the biochemistry and biophysics platform and animal genotyping facility of Neurocentre Magendie (INSERM). The microscopy was done at the Bordeaux Imaging Centre, a service unit of CNRS-INSERM and Bordeaux University, a member of the national infrastructure France BioImaging supported by the French National Research Agency (grant ANR-10-INBS-04).

## Author contributions

Experimental conception and design: D.C., C.H. and Y.H. Manuscript preparation: Y.H. All authors discussed the results and edited the manuscript. Behavioral experiments: H.E.O., C-L.Z., U.F., C.C. and A.L.-S.-A. Electrophysiology: U.F., F.L., P.M., L.V. and Y.H. Animal surgeries: H.E.O., C-L.Z., A.L.-S.- A., U.F., and A.N. Data analysis: M.L. and C.D. Competing interests: The authors declare that they have no competing interests. All reagents are available upon request. The mouse line can be provided pending a completed material transfer agreement. Requests for all materials should be submitted to Y.H. at.

## Materials and Methods

### 1. Biological models

Experiments in this manuscript were conducted on 6 to 12 weeks old male mice belonging to two strains: C57BL6/J wild type and C57BL6/J transgenic AP-GluA2 knock-In (KI, maintained on a C57BL6/J background) strains. Mice were kept on a 12-hour light/dark cycle and provided with *ad libitum* food and water, except for food restriction associated with behavioural testing (see below).

Mice were housed with 3-5 littermates except when demanded by the protocol. The experimental design and all procedures were in accordance with the European Guide for the care and use of laboratory animals. All procedures were validated by the ethical committee of animal experimental of Bordeaux Universities (CE50; Animal Facility PIV-EXP, APAFIS#18507-201901118522837; Animal Facility A1, APAFIS#4552 2016031019009163; Animal Facilities Neurocentre Magendie and PIV-EXPE, APAFIS#13515- 2018021314415739).

AP-GluA2 KI strain was developed and validated as a mouse model for AMPA receptors mobility^1^i^5^n. AP-GluA2 KI mice are similar to wild type C57BL6/J mice in terms of weight, size, growth or fertility, but also for tested cognitive abilities ^15^. At the genetic level, this strain presents a substitution of the endogenous GluA2 subunit of the AMPA receptor by a genetically modified one bearing an AP-(Acceptor Peptide) tag on the extracellular domain of the subunit. In the presence of the BirA ligase enzyme, which is not endogenously expressed, AP can bind Biotin, yet present in the murine brain. Thus, expression of AMPA receptors bearing biotinylated GluA2 subunits is restricted to neurons in which BirA ligase has been introduced by viral transfection, allowing targeting of AMPAR cross- linking. Indeed, presence of extracellular tetrameric Neutravidin consecutive of intracranial administration leads to immobilization of AMPA receptors at the synaptic and peri-synaptic space (see^15^).

### 2. Surgery

Various surgery protocols were performed depending on the aim of the procedure, dividing into two major subgroups: stereotaxic injections and chronic implantations. They eventually shared some common steps, hereby listed. Surgery protocol were similarly applied for both mouse strains.

#### ***i-*** Common surgery procedures

Mice were anaesthetised through exposure to the anaesthetic gas agent, Isoflurane (4% mixed with air) for 4 minutes and anaesthesia was maintained all throughout the surgery at 2% mixed with air. Mice were positioned in the stereotaxic apparatus (David Kopf Instruments) on a heating pad and received a subcutaneous injection of Buprenorphine (100μl, 0.1mg/Kg) and a local injection of Lidocaine (100μl, 0.4mg/kg) for analgesia. The scalp was rinsed with Betadine to prevent infections. After incision and opening of the scalp, Bregma and Lambda point were identified in order to identify the region of interest using atlas coordinates (Paxinos). Finally, sutures were applied to close the incision point, and mice were subcutaneously injected with analgesic agent (Carprofen, 100µl, 4mg/kg), fed with powdered-nutrient enriched food and left recovering inside a recovery cage positioned on a heating pad for 30 to 120 minutes. A post-surgery care routine was observed for 2-6 days following the surgery, during which the weight and general shape of mice were monitored and analgesic drugs were administered if needed.

#### ***ii-*** Stereotaxic injection

Stereotaxic injection was performed on mice aged 6-10 weeks. Injected viral vectors were charged inside 10μl graduated glass Hirschman pipets (ref. 960 01 05, Germany) and pressurized via a pressurized via a 5ml syringe (Terumo). The pipette was automatically descended into the target region at a speed of approximately 20µm/s (3 injections sites in the dorsal CA3; coordinates: AP - 2.35; ML ±2.65; DV -2.5/-2.0/-1.5). Injection of 250nl of the product was performed manually by applying low but constant pressure on the syringe. The pipette was maintained in position within the target region for 5 minutes after the end of the injection to allow local diffusion of the injected product, then retracted at a slow speed. When combined with other surgical procedures, stereotaxic injection always preceded stereotaxic implantation.

#### ***iii-*** Chronic implantation

Chronic implantation of guide cannulas and/or electrodes was performed on mice aged 6 to 10 weeks. Various types of implants were used, all of which are detailed in the dedicated “Implanted materials” section, but surgical procedures were common to all. Prior to proper implantation, the skull was prepared by briefly – 3 to 5 seconds - applying Peroxidase RED ACTIVATOR (Super-Bond, Sun Medical Co) to remove the periosteum. Single guide cannulas were manually descended into the target region at a speed of approximately 20µm/s (coordinates CA1: AP -1.95; ML ±2.25; DV -0.55; angle 30°; coordinates CA3: AP -2.35; ML ±2.65; DV -1.2). Electrodes were descended into the target region using a micromanipulator at 1µm/s for the last third of the descent, to reduce tissue damage caused by the implantation (coordinates: AP -2.35; ML ±2.3; DV -1.7; glued guide cannulas coordinates: AP -2.35; ML ±2.65; DV -1.2). Guide cannulas and electrodes were fixed with dental cement (Super-Bond, Sun Medical Co). After implantation mice were housed alone to prevent implants’ damaging.

### 3. Implanted materials

#### ***i-*** Guide cannulas

We used stainless steel guide cannulas (Bilaney 26 gauge, 1.5mm of length; PlasticOnes). Prior to the surgery, guide cannulas were kept in alcohol to minimize the risk for bacterial contamination and plugs were maintained on them at all time to avoid penetration of external material. When intracranial injections had to be combined with extracellular field recording, Guides were glued directly on the electrode connector and were obstructed with a metallic dummy cannula to avoid penetration of external material. Intracranial injection was performed on awake mice, either loosely held in the manipulator hands (for short injections) or free to move inside their home-cage.

Injections were performed through injection cannulas (Internal Cannula FIS 2.5mm guide, Bilaney; 0.5mm projection) and via an automatic pump (Legato 101, Kd Scientific Inc.) that applied a constant pressure using 1µl Hamilton syringes (7101 KH), allowing the regulation of injection speed (antibodies: 100nl/min; Neutravidin: 50nl/min). Pre-Learning and pre-test Injections were performed 1 hour before the beginning of the behavioural protocol; pre-rest injections were performed immediately after the last session of the behavioural protocol.

#### ***ii-*** Recording Electrodes

For hippocampal ripples recordings, bundles of Nichrome wires (diameter: 13µm, Sandvik Kantal) were connected to an 18 male connector (nano 18 positions 2 guides ISC-DISTREL SA Omnetics) were passed through a guide cannula (see **supplementary figure 2**) to protect them from damage and spreading while entering into the brain.

### 4. Chemicals

#### ***i-*** Viral vectors

All viral vectors used for the experiments described in the Results section are Adenoviruses and their engineering is detailed in Getz et al. 2021. Ongoing production was assured either by the viral core facility of the Bordeaux Neurocampus IMN or by Charité Universitats medzin Berlin or viral vectors were ordered on Addgene. All virus were stocked at -80°C for long-term storage, conserved at 4°C during surgery preparation and injected at room temperature. 250nl per injection site of viral vector solution were administered through stereotaxic injection during surgery.

- **pAAV9a-pSyn-BirA-ER-IRES-eGFP** (5.6 x 10^13 gcp/ml, IMN). The pSyn promoter allows the expression of the BirA enzyme in all neuronal types without distinction. BirA ligase expression promotes biotinylation of the extracellular portion of the GluA2 subunit of AMPA receptors, thus inducing AMPA receptors cross-linking in the presence of Neutravidin. eGFP is used as a tag do identify neurons expression the enzyme.

- **pAAV9a-pSyn-IRES-eGFP** (1.8 x 10^13 gcp/ml, IMN). Lack of BirA ligase coding sequence makes this viral vector a control for un-catalysed Biotin binding to GluA2 subunits bearing the AP.

#### ii-Antibodies

Production and conservation of antibodies used for the experiments detailed in the Results section of this manuscript is detailed in ^14^. All antibodies were stored at -80°C for long-term storage, conserved at 4°C for maximum 1week preceding injection and injected at room temperature. 500nl per injection site of antibody solution were administered via intracranial injection in the awake, freely moving mouse.

- **Antibody against GluA2 subunit of AMPA receptors (clone: 15F1)** (2.9mg/ml). This antibody is a monoclonal divalent IgG-κ directed against the extracellular domain of GluA2 subunit of

AMPA receptors. The divalent nature of this antibody allows for binding of 2 target GluA2 subunits at the same time, therefore promoting AMPA receptors cross-linking. In vitro, a washout time of 8 hours due to internalization of clustered receptors has been observed.

- **Fragment Antigen-Binding (Fab)** (2.9mg/ml). The antigen-binding portion of the antibody directed against GluA2 subunits was isolated and used as monovalent control for cross-linking.

- **Antibody against GFP** (2.9mg/ml). This antibody is a divalent IgG-κ from murine clones 7.1 and

13.1 (11814450001, Roche). As murine neurons do not physiologically synthetize GFP, this antibody was used as control for unspecific antibody binding.

#### iii-Others

- **Neutravidin** or **NA.** Texas Red-conjugated Neutravidin (8.33µM; Invitrogen, A2665) was used to operate cross-linking of AMPA receptors in the AP-GluA2 KI mouse model. 500nl per injection site of Neutravidin solution were administered via intracranial injection in the awake, freely moving mouse.

- **monomeric Streptavidin** or **mSA** was produced and conjugated to STAR 635P (Abberior,ST635P) using N-hydroxysuccinimide ester–activated fluorophore coupling as previously described.

500nl per injection site of mSA solution (concentration: 8.33µM) were administered via intracranial injection in the awake, freely moving mouse.

- **Saline Physiological solution** . Saline physiological solution was used as control for cross-linking in the AP-GluA2 KI mouse model. 500nl per injection site of Saline solution were administered via intracranial injection in the awake, freely moving mouse.

### 5. Behavioural protocol

Delayed Spatial Alternation (DSA) task. The DSA task is a delayed non-matching-to-place task used to assess special navigation and cognitive functions in rodents ^16^.

#### ***i-*** Food restriction

Food restriction was required to assure mice’s motivation. Mice were weighted right before food withdrawal and this weight was used to calculate the 85% of weight-loss limit that was fixed for protocol termination. On the first day of restriction, mice were fed with Perles pasta of the same type as those that were used to bait the maze during the behavioural task, in order to habituate them to the new food. On subsequent days, mice were fed at the end of all behavioural manipulation with 2-3g of powdered nutrients-enriched food, in order to maintain them to about 85% of their initial weight.

#### ii-Materials

A custom-made, semi-transparent white P V C Y m a z e w a s u s e d t o t h e t a s k . A l l t h r e e a r m s a r e identical (40cm length, 8cm width, 15cm high walls), except for an additional closable rectangular chamber (15cmx25cm) bridged to the “Starting arm”. Arms are spaced by a 120° angle. An opaque small container was positioned at the end of each “Goal arm” to serve as food well for reward delivery. Environmental cues are positioned on the room walls surrounding the maze. Video recordings are realized through an infrared camera (Basler USB camera - ac1920-155um - Noldus) positioned on the ceiling upon the centre of the maze. The DSA tests were realised in conditions of dim light.

#### iii-behavioural paradigm

- **Habituation** . Habituation lasted for 5-8 days, depending on each individual mouse, and was divided into 3 phases. A first phase, starting before food restriction, consisted in 2-3 days of handling in order to habituate the mouse to be manipulated, especially for injection cannulas insertion and/or electrodes plug-in. Proper habituation for the task started on the second day of food restriction and consisted in multiple sessions of free exploration of the maze. The sessions were stopped when the mouse had eaten food-pellet in all three baited arms. The last phase of habi consisted in a single trial in which the mouse was positioned inside the “starting arm” (defined by position with respect to environmental visual cues) and had to collect a reward food-pellet in each of the two “goal arms”, with a time-limit of 1 minute.

- **Task.** The DSA task consists in 10 trials in which the left and right goal arms are alternatively baited with a rewarding chocolate pellet. During the first trial, the choice is forced toward the baited arm, setting the pattern of alternation (i.e. the reward-zone of the un-baited arm is made inaccessible through positioning a PVC slide at the entrance of the proximal portion of the arm; each consecutive session alternatively starts with a forced right or left choice). The 9 following trials rely on the mouse free choice of one of the two arms. A single trial can be repeated up to 6 times (“runs”) if the mouse makes consecutive mistakes, of which the sixth consists in a forced run in the baited arm direction. Once the mouse has reached the reward-zone of the chosen arm, access to the reward-zone of the unchosen one is restricted and the mouse is let spontaneously come back to the distal portion of the starting arm. After every run, the mouse is placed back in his home-cage and a delay of 30 seconds is respected before the mouse is allowed to explore the maze again.

During this delay period, the maze is cleaned with ethanol to prevent odour-based navigation. On the first day of training, 4 sessions are conducted, spaced by 30-60 minutes rest period; on the 2 following days, 2 sessions per day are realized, spaced by 30 minutes (see **Figure 1**).

Behavioural training was conducted between 8a.m. and 1p.m.

### 6. *In vivo* electrophysiological recording

#### i- Recording sessions

Electrophysiological recordings are realized by plugging a headstage (INTAN) containing 16 unity-gain operational amplifiers to each connector. Recordings were realised through the recording system OpenEphys. Recording were performed during resting periods in the mouse home cage positioned in an isolated closed box allowing cables suspension and infrared recordings by the camera (Basler USB camera - ac1920-155um - Noldus). A baseline is recorded for 3 hours long during a rest session the day preceeding the first training sessions. This recording is repeated the following day after the 4 sessions of tranings. The starts averagely 10 minutes after the end of the last habituation/training session of the day.

#### ***ii-*** LFP processing and Sharp Wave-Ripples detection

Electrophysiological data were imported on Matlab and down-sampled to 1kHz for storage and analysis speed convenience. Ripples detection was performed on Matlab scripts originally developed by Cyril Dejean. At first, referencing and band-pass filtering at 50 Hz eliminates noise oscillations common to all channels. Then, 100-250Hz band-pass filtering is used to detect ripple events that are selected if respecting following criteria: 1) its amplitude is higher than 5 standard deviations of the mean band-passed trace, 2) the event must be at least 30ms long and 3) two ripples must be separated by an interval of at least 45ms. Ripple’s characteristics are then computed, including: timestamp of the peak, intrinsic frequency, number of oscillations, mean amplitude (both on the filtered and the integrated trace), area under the integrated curve, duration (total and of each part preceding and following the peak), half prominence.

### 7. In situ slice recordings

#### ***i-*** Preparation of hippocampal slices

Mice males WT C57bl6/J were used at the age of 4 to 9 weeks. The extracellular ACSF (Artificial <u>C</u>erebro-<u>S</u>pinal <u>F</u>luid) solution utilized for slice recordings is composed of: 119 mM NaCl; 2.5mM KCl; 1.3mM MgCl2; 2.5mM CaCl2; 10mM glucose; 1mM NaH2PO4; 26mM NaHCO3. The cutting solution is an ice-cold sucrose solution (1-4°C) composed of 2 mM KCl; 2.6mM NaHCO3; 1.15mM NaH2PO4; 10mM glucose; 120mM sucrose; 0.2 mM CaCl2 and 6 mM MgCl2. Both solutions are oxygenated with carbogen (95% O2, 5% CO2, pH 7.4 at 37°C, 290-310 mOsm/L). For brain dissection, mice are anesthetized with 5% isoflurane for 2 min before decapitation. The head is immersed in the iced sucrose solution. The removed brain is immersed for 4 minutes in iced oxygenated sucrose solution and then placed on a cellulose nitrate membrane to separate and position the hemispheres in the vibratome (Leica VT1200s) to obtain 400 µm horizontal slices (speed of 0.1 mm/s). Once produced, slices are semi-immersed in a dedicated incubation chamber, oxygenated and maintained at 35°C in a water bath for at least 2 hours before starting the recordings.

#### ***ii-*** In vitro electrophysiological recordings

Recordings are made in an S-shaped recording chamber, maximizing oxygenation while preventing slice movement caused by the 3.5 mL/min perfusion flow. Field recordings are obtained using glass micro pipettes stretched with a PC-10 (Narishige, Japan) and broken at their tip to decrease the resistance (<0.5 M!l). Depending on the experimental configuration, the pipette is filled either with ACSF or supplemented with antibodies, IgG IZ-GluA2, 15F1 (IgG) or IgG Fab (Fab).

Electrophysiological recordings are obtained by a MultiClamp 700B (Molecular Devices, Foster City, CA) using Clampfit software (Molecular Devices, Foster City, CA). Electrical stimulation is provided by a CBCSE75 concentric bipolar electrode (FHC, Phymep, France) and an external A.M.P.I Iso-flex stimulator (Scop Pro, France). Synaptic field are recorded in the stratum radiata (sr) of CA1 and CA3 to measure evoked fEPSPs, and to test the presence of propagated SPW-Rs. Depending on the configuration, stimulation electrodes are placed in the stratum radiatum of CA1 or CA3 to stimulate respectively CA3➔CA1 and CA3➔CA3 axons and elicit basal (0,1 Hz) or high frequency stimulations (HFS, 100 Hz, 1 sec train repeated 3 times each 30 sec) to induce long term potentiation (LTP). Wen stimulating CA3 sr, a train of 10 stimulations at 1 Hz is first applied to test for eventual contamination by DG mossy fibers that display frequency facilitation.

### 8. Perfusion and Histology

Mice were anaesthetized with a mix of Kétamine and Xylazine (100mg/20mg/kg) diluted in a saline solution; 10µl of solution per gram weighted by the animal were administered via peritoneal injection. Perfusion with Paraformaldehyde (PFA, concentration: 4%) was realised on the anaesthetized mouse and brain were initially collected without being evicted from the scalp. After 48h of storage in PFA at 4°C temperature, implants and scalp were removed and the brain was washed three times in PBS solution (concentration: 1%) and then stored in PBS for 24-72 hours at 4°C. Slicing was performed with a vibratome (Leica VT1200s). Coronal slices of 60µm thickness were collected at a speed of 30-50mm/s from the regions of interest and stored in PBS for 24h before being mounted on slides and covered with Fluoromont-G (complemented with DAPI for cellular nuclei staining, Thermofisher Scientific).

Image acquisition of slides was performed with an epifluorescence microscope (Nikon Eclipse NI-U) coupled to an illumination system (Intensilight C-HGFI, Nikon) and a camera (Zyla sCMOS, Andor Technology, Oxford Instruments).

---

**Sup. Figure 1:**
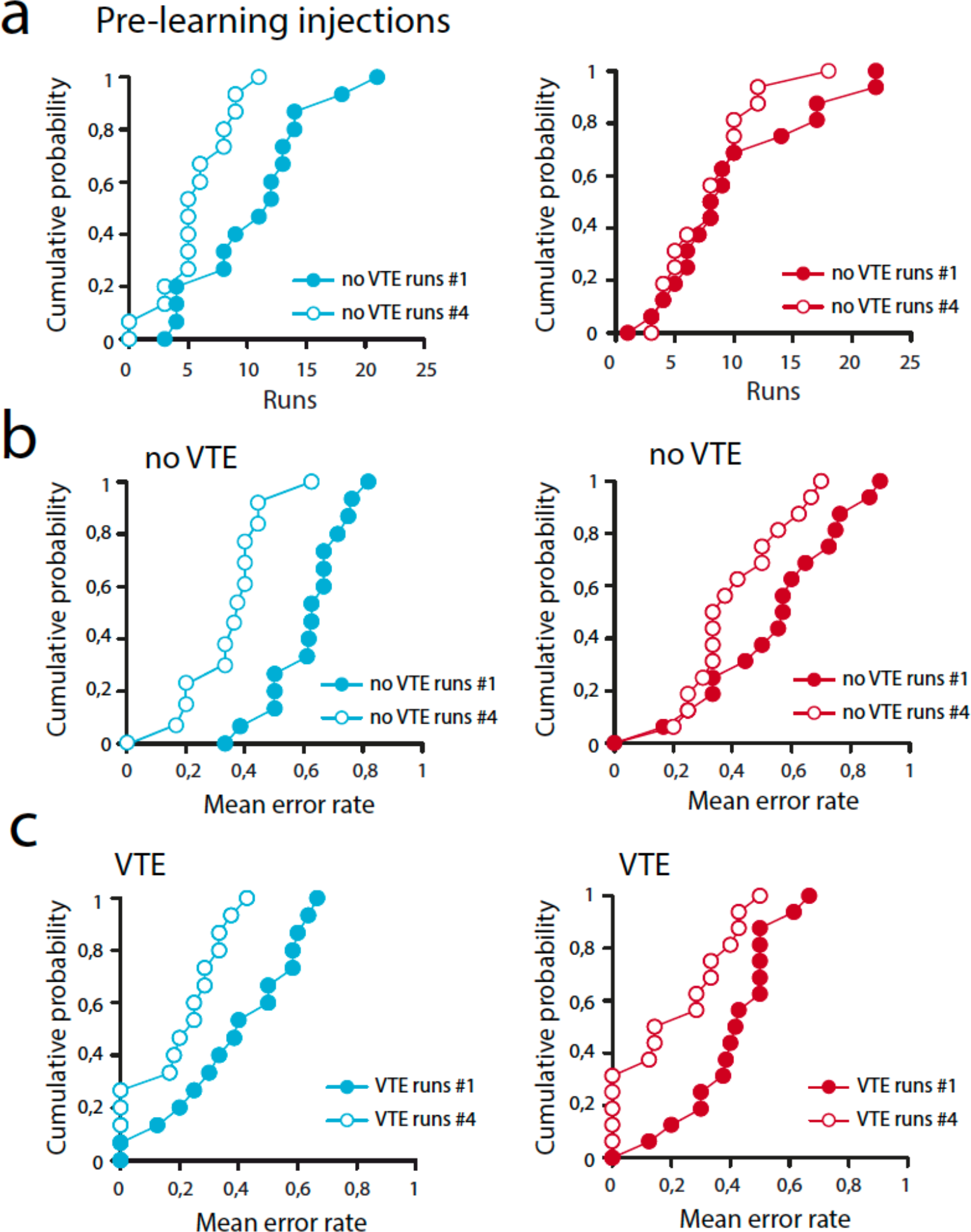
dHPC AMPAR X-linking preserve DSA rule encoding. Same presentation as Figure 2. **a**: Cumulative single animal data for no VTE run numbers in session session#1 and session session#4. **b-c**: Choice accuracy at VTE and no VTE run numbers in session session#1 and session session#4.

**Sup. Figure 2:**
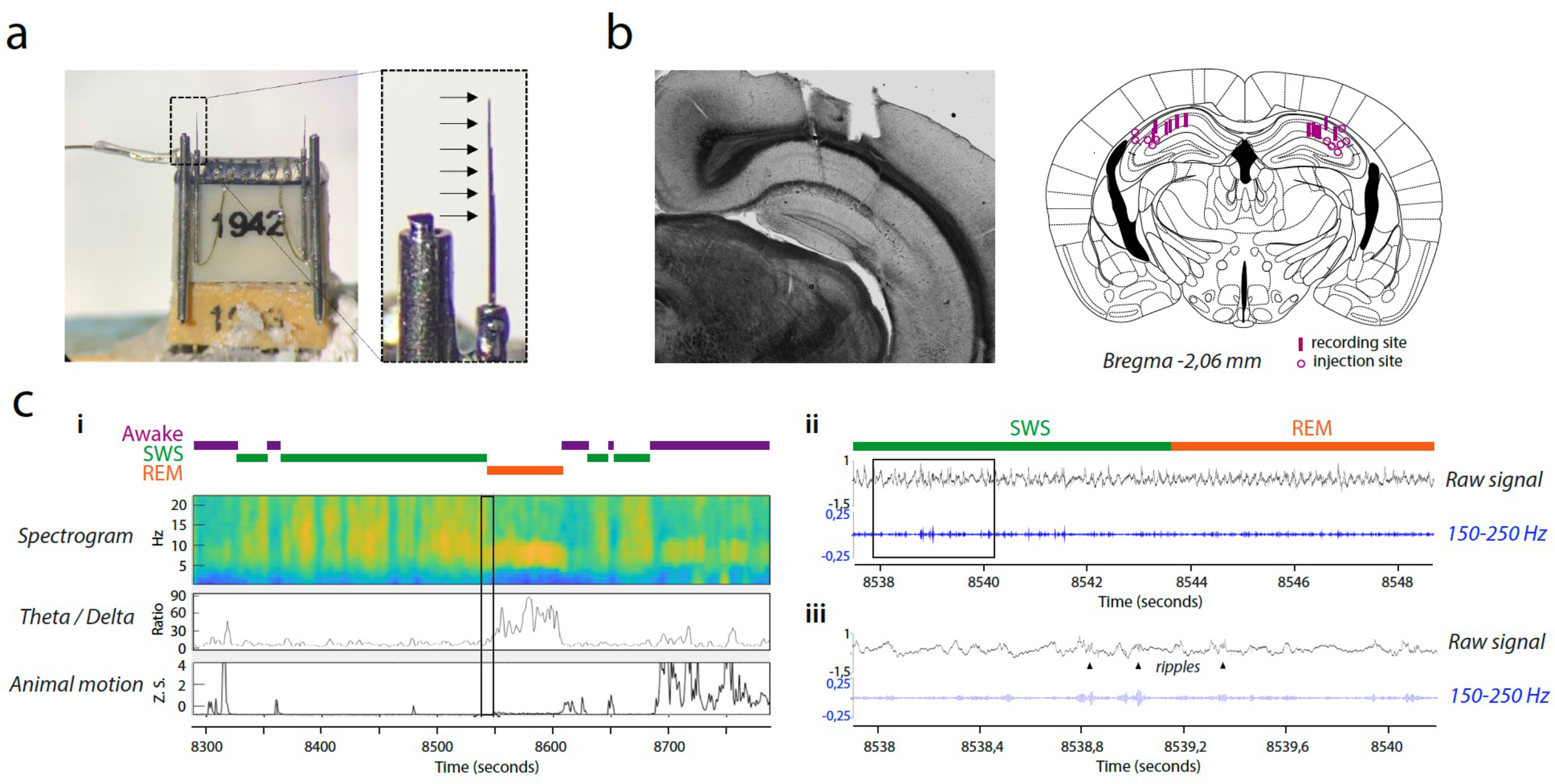
Combined dHPC electrophysiological and pharmacological experiments. **a**: photograph of the design implant. Ripple recordings are performed *via* 2x6 wires cut at 200 micrometres intervals in order to target the stratum pyramidale of the CA3 region. Lateral to each bundle, are positioned cannulas that will be used for drug injection. **b**: micrograph of coronal slice of implanted mouse brain. Arrows identified the bundle of recording wires, and the tip of the implanted cannula. To perform injection, an injection cannula projects out of 1 millimetre within the hippocampus and is retracted back during the injection in order to cover a large portion of the dHPC. **c**: Example of injection of anti-GluA2 antibody that is detected by immunofluorescence (see methods for more details). **d**: Analytical pipeline for sleep and ripple detection. Animal tracking (motion) allows separating awake and sleeping states. Within sleep periods, LFP analysis of the Theta/Delta range separate REM (high theta/low delta) and SWS (low theta/high delta) phases. **e**: In parallel, LFP are band pass filtered between 150-250 Hz in order to extract ripples oscillations (see methods). A zoom on detected ripples is shown in **f** (triangles).

**Sup. Figure 3:**
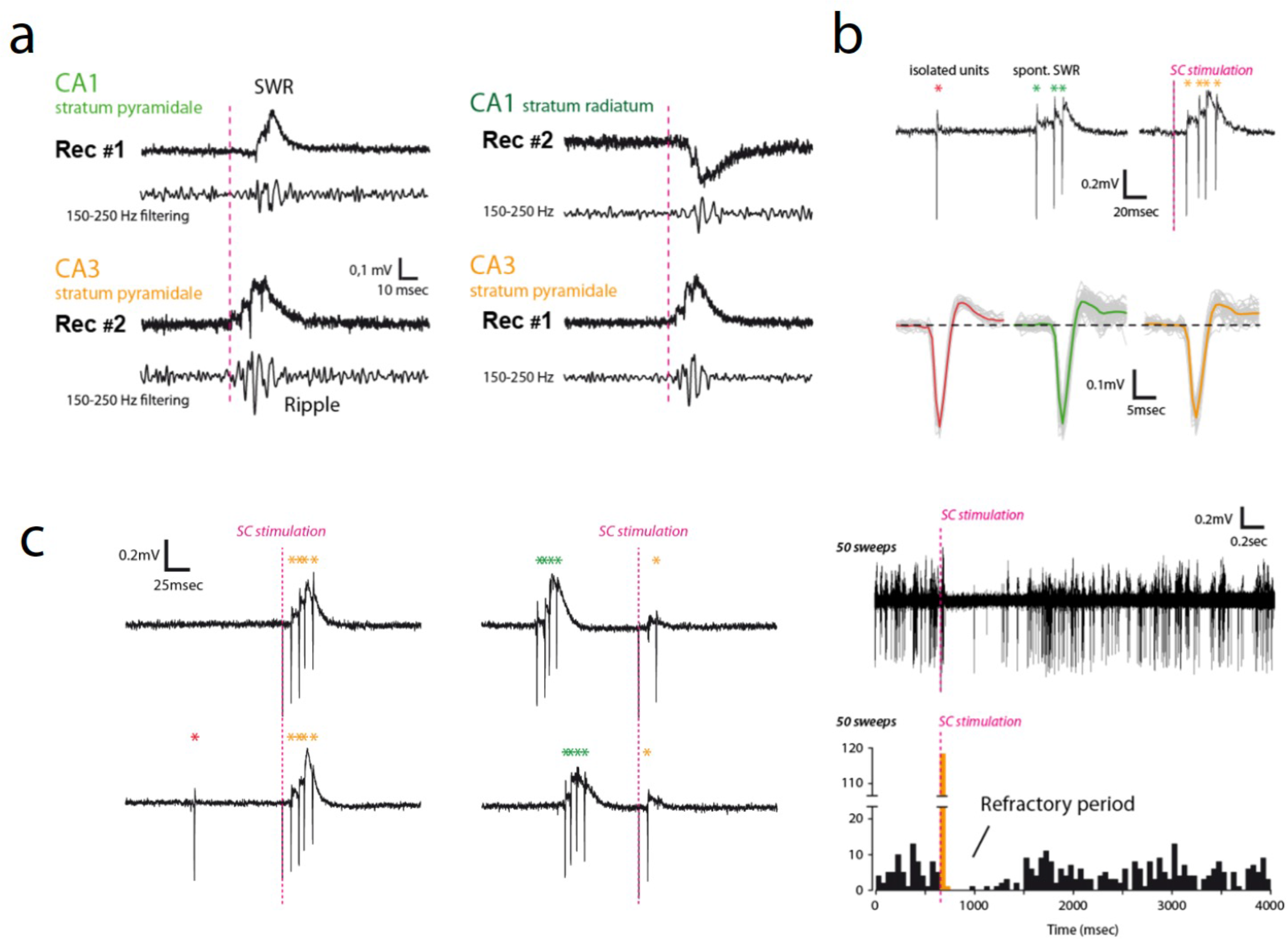
Characteristics of SWRs recorded in hippocampal acute slices. **a**: SPW-Rs events recorded in acute hippocampal slice are composed of dendritic synaptic wave that polarity is changed depending on electrode position (upward in CA1-sp (left), downward in CA1-sr (right)). Filtering at 150-250 Hz shows high frequency oscillations (ripples) associated to the wave. Note that a significant delay exists between CA3 SWRs and those recorded in CA1 area, suggesting that they are occurring first in the CA3 region. **b**: Typical example of a CA3-sp single unit recording showing that spontaneous or evoked (through CA3 axons stimulations) SWRs are indeed leading to focal neuronal activation. Bottom: waveforms of the single unit in the three conditions (alone, spontaneous SWRs and evoked SWRs). **c**: interaction between spontaneous and evoked SWRs. Typical example of refractory period observed for consecutive SWRs that probably limit their activity in acute slices. Left: examples of collision between spontaneous and evoked SWRs. Right: No spontaneous SWR is observed after the generation of an evoked SWR, suggesting that these oscillations may require time-dependent regenerative mechanisms.

**Sup. Figure 4:**
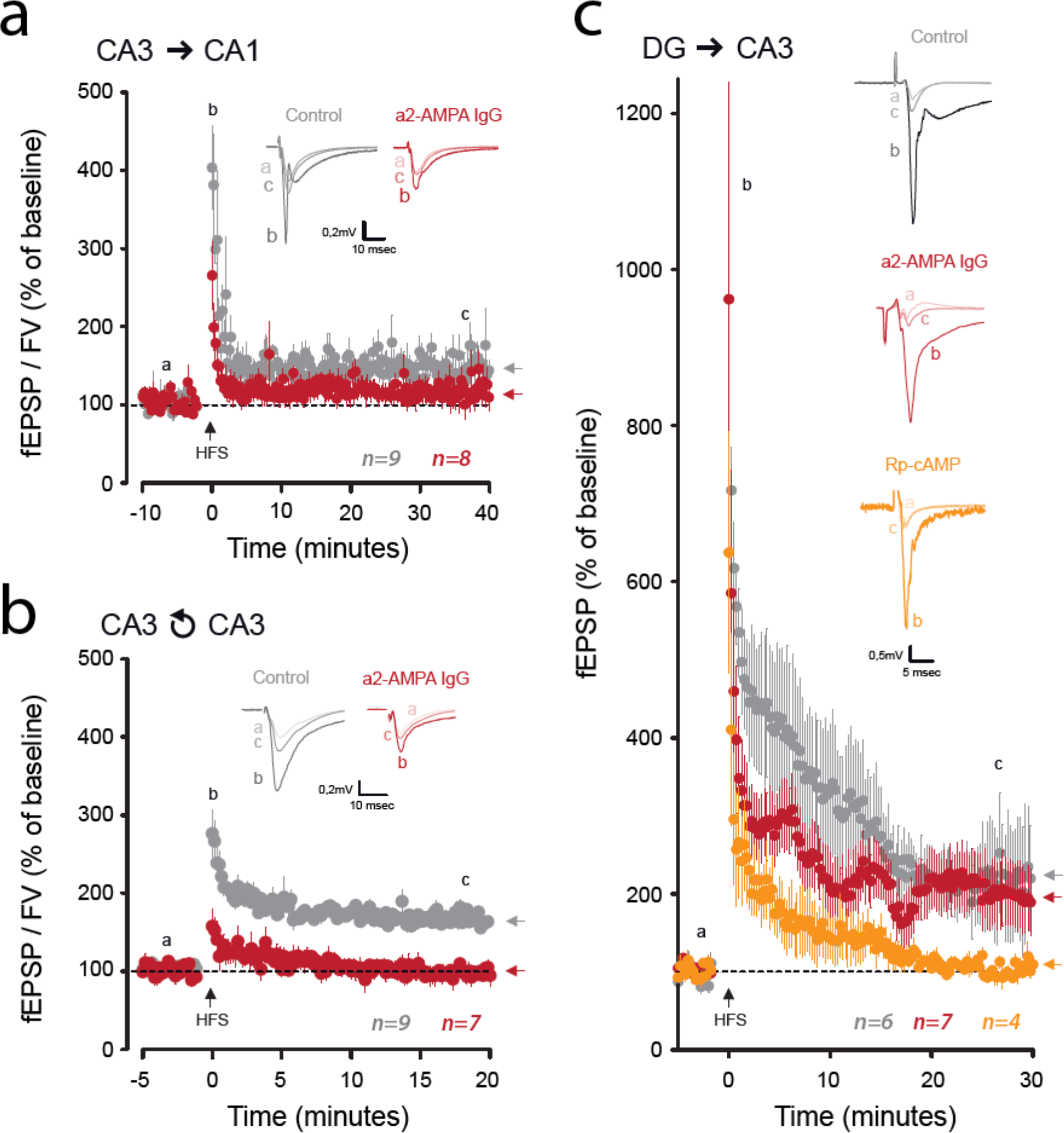
AMPAR X-linking impair LTP at CA3 **➔**CA1, CA3**➔**CA3 but not DG**➔**CA3 synapses. **a**: Time course of fEPSP/FV slopes ratio before and after high frequency stimulations (HFS) in control (grey) and AMPAR X-linking conditions (red). Note the absence of potentiation in presence of anti GluA2 IgGs (arrows). In insert, representative traces obtained during baseline (a), immediately after HFS (b) and 35 minutes after HFS (c) in both conditions. **b**: Same presentation as in **a**. HFS stimulation was applied within the CA3 *stratum radiatum* to isolate CA3 ➔CA3 recurrent synapses. Note the absence of potentiation in presence of anti GluA2 IgGs (arrows). In insert, representative traces obtained during baseline (a), immediately after HFS (b) and 20 minutes after HFS (c) in both conditions. **c**: Same presentation as in **a**. HFS stimulation was applied within the CA3 *stratum lucidum* to isolate DG➔CA3 synapses. Note the potentiation that is still present in the anti GluA2 IgGs condition, but not when slice was preincubated with Rp-cAMP to block PKA activity (arrows). In insert, representative traces obtained during baseline (a), immediately after HFS (b) and 20 minutes after HFS (c) in all conditions.

**Sup. Figure 5:**
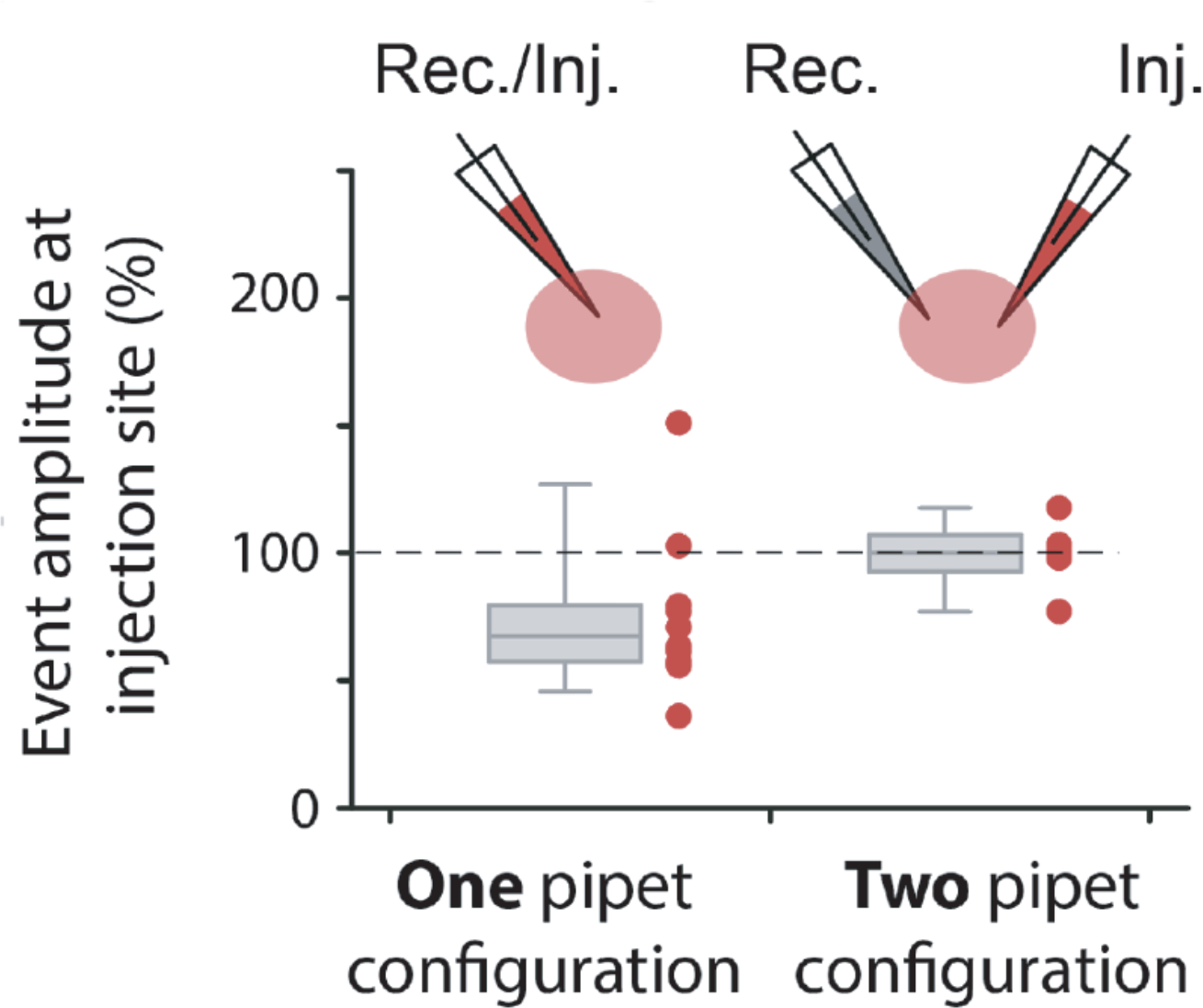
AMPAR X-linking effect on SWR amplitude is due to injection procedure. We compared the local effects of anti-GluA2 antibodies on SWR amplitude in experiments in which it was introduced via the recording pipet (One pipet configuration) or using another pipet than the recording one (two pipet configuration). As can be seen, the decrease in amplitude of SWRs was due to the positive pressure applied in the pipet that probably moved away the tissue locally. Thus we conclude that, as previously observed ^14^ that AMPAR X-linking was not affecting basal transmission, and thus leave spontaneous SWRs unaffected.

## Notes

### Competing Interest Statement

The authors have declared no competing interest.

